# Personalized Neuroimaging Reveals the Impact of Children’s Interests on Language Processing in the Brain

**DOI:** 10.1101/2023.03.21.533695

**Authors:** Halie A. Olson, Kristina T. Johnson, Shruti Nishith, Isabelle R. Frosch, John D.E. Gabrieli, Anila M. D’Mello

## Abstract

Cognition is shaped by individual experiences and interests. However, to study cognition in the brain, researchers typically use generic stimuli that are the same across all individuals. Language, in particular, is animated and motivated by several highly personal factors that are typically not accounted for in neuroimaging study designs, such as “interest” in a topic. Due to its inherently personal and idiosyncratic nature, it is unknown how interest in a topic modulates language processing in the brain. We conducted functional magnetic resonance imaging (fMRI) in 20 children (ages 6.98-12.01 years, mean(SD)=9.35(1.52), 5 female/15 male) as they listened to personalized narratives about a topic of specific interest, as well as to non-personalized generic narratives. We found that personalized narratives about a topic of interest increased activation in canonical language areas, as well as in reward and self-reference regions. Strikingly, we found that activation patterns elicited by topics of personal interest were more consistent across children, despite their idiosyncratic nature, than activation patterns elicited by narratives about an identical generic topic. These results reinforce the critical role that personal interests play in language processing in the human brain, and demonstrate the feasibility of using a personalized neuroimaging approach to study the effects of individually-varying factors such as interest in the brain.

## INTRODUCTION

Certain cognitive experiences are highly specific to the individual. However, the scientific study of these experiences largely ignores their personal nature, instead relying on experimental stimuli that are generic and identical across individual participants. In particular, language usage and experience – such as what we speak about and are motivated to listen to – are animated by several highly personal factors that are not often accounted for or considered in study design, including *interest*.

Here, we refer to an “interest” in a topic as the sustained fascination one has with a subject. Interests are inherently personal and idiosyncratic, marked by heightened focus and a lasting inclination to revisit the subject repeatedly (Harackiewicz et al., 2016; Renninger et al., 2014; Renninger & Hidi, 2015), and they can be powerful motivators of behavior. For instance, interesting materials have been used to motivate learning and improve academic achievement (Reber et al., 2018; Walkington & Bernacki, 2014), and children who score poorly on tests of reading comprehension for generic material often score better when these tests involve interesting (Shnayer, 1968) or familiar (Recht & Leslie, 1988) material. Shared interests also promote social engagement and generous behaviors in children (Sparks et al., 2017). For populations that struggle with language and language-relevant skills, interests can be a particularly potent motivator. Prior work in highly successful professionals with dyslexia found that a common theme underyling their eventual reading success was a passionate interest in a topic (Fink, 1995). Similary, in children with autism – a condition characterized in part by language and social communication challenges – several clinical studies found that scaffolding communicative interactions and interventions around topics of personal interest enhanced communication skills and social interaction (Arunachalam et al., 2024; Baker et al., 1998; Boyd et al., 2007; Charlop-Christy & Haymes, 1998; Harrop et al., 2019; Lizon et al., 2023; Suskind, 2014).

Although interests impact language and communicative behaviors, few people have studied how they impact brain function, especially in the context of language processing. One roadblock in the study of interest is its inherently personal and idiosyncratic nature, which is difficult to incorporate into traditional neuroimaging approaches which prioritize experimental control. These approaches generally do not consider inter-individual variability in participants’ experience with or interest in the stimuli, even though differences in observers’ interpretations of stimuli have been shown to impact brain responses (Varrier & Finn, 2022). For example, in studies of language, certain commonly-used linguistic metrics for matching stimuli (e.g., word frequency) can be misleading because they assume a common experience across individuals. While low-frequency specialized words like “Pikachu” or “Pokéball” may appear infrequently in general language use, they may be highly familiar to a child with a special interest in Pokémon. One way to incorporate individual variability with regards to interest in experimental stimuli is to *personalize* experimental stimuli within the context of a standard paradigm. Personalization can account for individual differences in experience, such as the word frequency example above, that cannot be addressed in other ways (e.g., matching on stimuli-level features, collecting huge sample sizes, using large corpora to calculate word frequency, etc.). This approach has been successfully applied in a handful of studies based on the intuition that personalization (e.g., a favorite food (Tomova et al., 2020), a video of a specific memory (Bainbridge & Baker, 2022), or a personal narrative story to mimic internal thought (Kim et al., 2024)) might be the most effective and ecologically valid way to elicit or decode neural responses in brain regions that process those stimuli.

Here, we apply a personalized neuroimaging approach to examine the effect of children’s interests on language processing in the brain by creating *individualized* language stimuli specific to each child’s interest. This approach allowed us to capture the personalized nature of interest within the context of a standard paradigm across all participants, examine the effects of interest on the brain, and empirically assess the feasibility of using a personalized neuroimaging approach to study this highly individual factor.

## MATERIALS AND METHODS

### Participants

Data were analyzed from n=20 children (ages 6.98-12.01 years, mean(SD)=9.35(1.52), 5 female/15 male). All children were native speakers of English, had normal or corrected-to-normal hearing and vision, had no contraindications for MRI (e.g., metal in the body), and had a qualifying special interest (see **Personal Interest Screening** below). Additional exclusion criteria included a diagnosis of major neurodevelopmental or psychiatric disorders and/or language difficulties. A total of 27 children were originally recruited for participation, but n=7 were not included in these analyses due to excessive motion for the language task (n=5), incidental findings (n=1), and incomplete data (n=1). Parents provided informed consent, and children provided assent to participate. This protocol was approved by the MIT Committee on the Use of Humans as Experimental Subjects.

### Personal Interest Screening

Parents or caregivers expressed interest in the study via an online screening survey. If a child was potentially eligible (i.e., appropriate age, no exclusions based on the criteria listed above, and parent-reported presence of a significant interest, hobby, passion, or affinity), a member of the research team conducted a phone screening and discussion with caregivers to confirm eligibility and to ask follow-up questions about the child’s interest. Criteria for the presence of a personal interest were as follows: (1) the child must engage with the interest for at least an hour per day on average (or would engage with that interest for the specified amount of time if there were restrictions in place, such as screen time limits), (2) the child must have had the same interest for at least the last two weeks, and (3) the child must have an interest that could be represented by videos. Caregivers, in collaboration with their children, were then asked to provide video clips pertaining to their child’s interest, which we used as inspiration for the personalized narratives in the fMRI experiment (see **Personalized Stimuli Creation**).

### Experimental Design

#### Experimental Protocol

Participants completed 1-2 study sessions, which involved behavioral testing and a neuroimaging session. The neuroimaging session included an anatomical scan, a functional run of a task which involved watching the participants’ selected interest videos and nature videos (see **Supplementary Materials**), a functional run of the personal interest language task (audio only; see **Functional Personal Interest Narrative Task** for details below), and optional additional scans that varied between participants. These options included a resting state scan, neural adaptation tasks involving faces, objects, and auditory words, a separate language task, a diffusion scan, and additional runs of the personal interest tasks. Various strategies were employed to prepare children for the neuroimaging session, including familiarization with a mock scanner and practice with task instructions outside the scanner.

#### Behavioral Measures

We conducted assessments of nonverbal reasoning and language skills to ensure all children could understand task procedures and instructions, control for cognitive reasoning (i.e., nonverbal IQ), and characterize language skills (**Table 1**). Nonverbal cognitive reasoning was assessed via the matrices subtest of the Kaufman Brief Intelligence Test, 2nd edition (KBIT-2; Kaufman & Kaufman, 2004). Language skills were assessed via the verbal composite score of the KBIT-2, including the vocabulary and riddles subtests. Parents completed a set of questionnaires about their child during the visit including questions regarding demographic and developmental histories (e.g., language onset).

**Table 1:** Demographics and interests of participants. Group means (and standard deviations) for several demographic and cognitive measures. KBIT = Kaufmann Brief Intelligence Test. Note that the average for KBIT verbal composite is missing one participant who did not complete the assessment.

|  |  |
| --- | --- |
| Age (years) | 9.35(1.52), range = 6.98 – 12.01 |
| KBIT Matrices (standard score) | 118.65(15.39), range = 87.00 – 140.00 |
| KBIT Verbal Composite (standard score) | 121.00(12.99), range = 98.00 – 142.00 |
| Interests | Soccer (n=3)<br>Baseball and football (n=2)<br>Basketball<br>Fishing<br>Fortnite<br>Minecraft (n=3)<br>Pokémon<br>Lego Marvel superheroes<br>Animals from PBS show<br>Baking shows<br>Musicals<br>Harry Potter<br>Art videos (n=2)<br>Transit Systems |

#### Non-Personalized “Neutral” Stimuli Creation

Seven video clips of different nature scenes were selected from an open-access online repository (YouTube). Based on these scenes, seven 16- second engaging audio narratives of the scenes were created, and a female experimenter (HAO) recorded the descriptions in a sound-proof booth. The audio files were trimmed to be exactly 16 seconds. All participants listened to the same set of non-personalized narratives but to a unique set of personalized narratives (see **Personalized Stimuli Creation** below). No children reported nature as a special interest.

#### Personalized Stimuli Creation

Caregivers, in collaboration with their child, provided links to online video clips (e.g., YouTube links) that captured their child’s personal interest, including timestamps for their child’s favorite parts of the videos. We cut seven 16-second clips from the provided videos (capturing each child’s favorite part of the videos if provided), and wrote short audio narratives based on the scenes from the selected video clips. The same female experimenter recorded the descriptions, following an identical protocol to the non-personalized stimuli. Language narratives were approximately matched between participants by avoiding personal pronouns, using simple vocabulary (allowing for interest-specific terms), and using short sentences. Due to the unique nature of the personal interests, the personal-interest narratives tended to have more specific nouns – e.g., “Alewife Station” or “Lionel Messi” – than the neutral narratives. Some, though not all, personally- interesting narratives also contained social content, such as discussions about specific people or human interactions; however, many personal interests were not social in nature (see **Table 1**). Both the personally-interesting and neutral narratives included action verbs and sensorially evocative descriptions. See OSF (https://osf.io/dh3wq/) for the neutral and personally-interesting narrative transcripts for all children included in this analysis.

Linguistic and paralinguistic features, including word count, words per sentence, syllables per word, number of sentences (all from https://goodcalculators.com/flesch-kincaid-calculator/), parts of speech (from the spaCy Python library; https://spacy.io/), word frequency (from the wordfreq Python library; https://pypi.org/project/wordfreq/) and emotional valence (using the TextBlob Python library to calculate sentiment polarity; https://textblob.readthedocs.io/en/dev/) measures, were analyzed across all narratives. Non-parametric pairwise comparisons were conducted using Wilcoxon rank-sum tests to identify specific differences between individual subjects’ narratives and the neutral narratives. After applying a Bonferroni correction for multiple comparisons (adjusted p<0.0025 for 20 comparisons), no significant differences were found for the number of sentences, syllables per sentence, emotional valence, adverb usage, verb usage, and adjective usage across narratives. Some subjects showed significant differences for words per sentence (n=1), word count (n=2), word frequency (n=3), number of nouns (n=7), and number of proper nouns (n=12). Because the NEUTRAL condition contained no proper nouns, we also combined nouns and proper nouns and then found no condition differences.

Box plots and raw outputs of the statistical analysis for all features across narratives are provided in the Supplementary Material.

#### Functional Personal Interest Tasks

Participants completed two separate personal interests tasks within the scanner: a video task wherein participants watched video clips of their personal interests and nature scenes (beyond the scope of the current manuscript, see details of task design and analyses in **Supplemenatry Materials)** and a narrative task. During the personal interest language task, participants were asked to passively listen to spoken narratives presented binaurally via MRI-compatible headphones using a block-design paradigm. The task consisted of three conditions: INTEREST, NEUTRAL, and BACKWARDS narratives. In the INTEREST condition, participants listened to the personalized narratives about their specific interests. In the NEUTRAL condition, participants listened to non-personalized narratives describing nature scenes. Nature content included in the neutral narratives was similarly familiar to all children and unrelated to any child’s personal interest. In the BACKWARDS condition, participants listened to incomprehensible reversed versions of the neutral narratives in order to account for lower-level auditory features of the narratives. Children listened to 7 narratives (16-seconds each) in each condition while viewing a static white fixation cross on a dark grey background. Each narrative was followed by an inter-stimulus rest block of 5 seconds (total of 21 narratives across three conditions and 22 rest blocks) cued by a grey fixation cross. To confirm that children were attending to the task without imposing significant physical or cognitive demands, we included a low-demand attentional check following each narrative. An image of a panda appeared on the screen directly after each narrative for 1.5 seconds, followed by a grey fixation and screen for 0.5 seconds. Children were instructed at the beginning of the study to press a button using their pointer finger via an MRI-compatible button box that they held in their hand every time they saw a picture of a panda. Task order was fixed across participants in an [ABCABC…] pattern: INTEREST, NEUTRAL, then BACKWARDS (see **Figure 1**).

**Figure 1:**
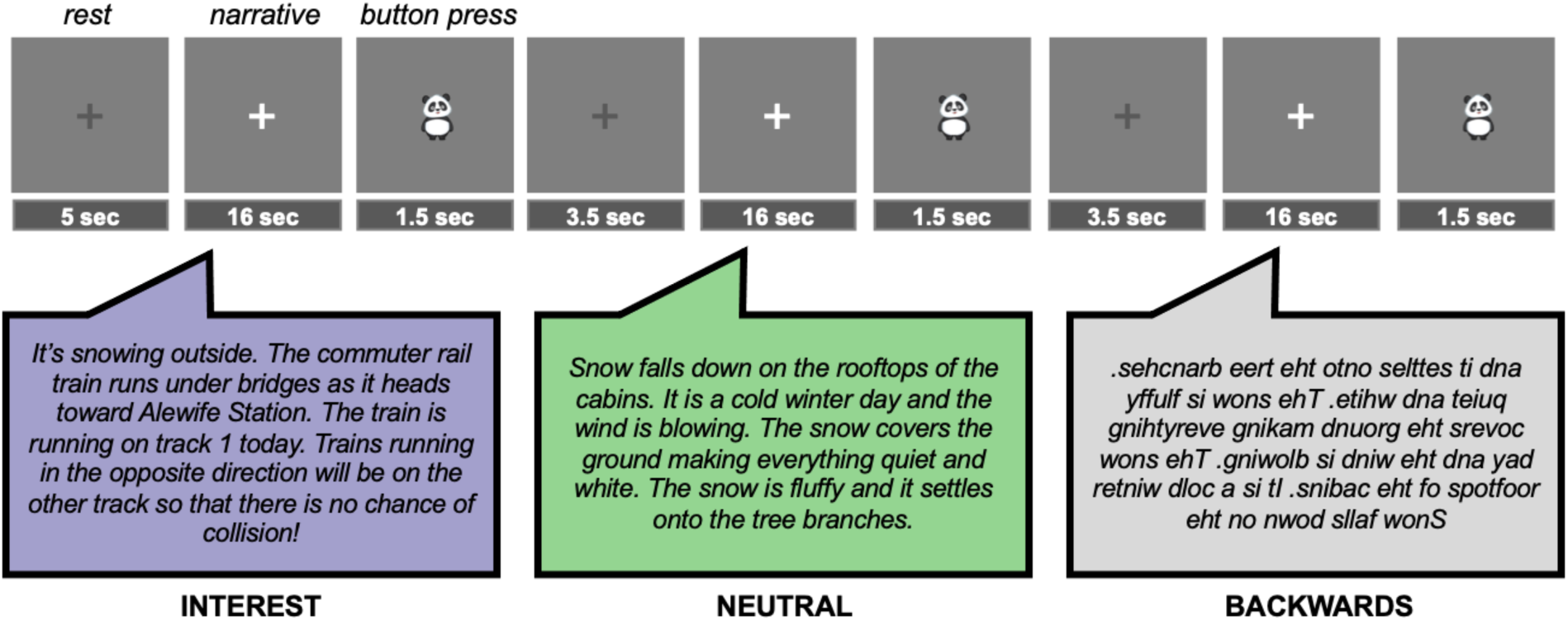
Task design. Participants passively listened to 16-second narratives in three conditions (INTEREST, NEUTRAL, and BACKWARDS) in a fixed order. Each run contained 21 blocks (7 per condition).

#### Acquisition

Data were acquired from a 3-Tesla Siemens Prisma scanner located at the Athinoula A. Martinos Imaging Center at the McGovern Institute at MIT, using a 32-channel head coil. T1-weighted structural images were acquired in 196 interleaved slices with 1.0mm isotropic voxels (MPRAGE; TA=5:08; TR=2530.0ms; FOV=256mm; GRAPPA parallel imaging, acceleration factor of 3).

Functional data were acquired with a gradient-echo EPI sequence sensitive to Blood Oxygenation Level Dependent (BOLD) contrast in 3.0mm isotropic voxels in 40 near-axial slices covering the whole brain (EPI factor=70; TR=2500ms; TE=30ms; flip angle=90 degrees; FOV=210mm; TA=7:47).

### Statistical Analysis

#### Overview of Analytical Approach

Our primary hypothesis was that personal interest would elicit stronger responses to linguistic stimuli in language regions. Thus, we first examined the magnitude of the response to each condition within canonical language regions (**Region of Interest Analyses**). To further explore the impact of interest on language regions’ function, we then examined functional connectivity between language regions. If personal interest content fundamentally changed the functional organization of the language network, then we would expect to see see different functional connectivity strength during the INTEREST condition compared to the NEUTRAL condition (**Task-based Functional Connectivity Analysis**). A second hypothesis was that the INTEREST condition would elicit higher activation in non-language regions relevant for processing interesting content relative to the NEUTRAL condition (such as reward regions). We therefore examined responses across the whole brain for the INTEREST>NEUTRAL contrast at both the group and individual level (**Whole Brain fMRI Analyses**). Finally, because we were interested in the feasibility and application of personalized stimuli in neuroimaging more broadly, we explored one potential *downside* of using idiosyncratic stimuli that vary across participants: the likelihood of greater between-particiant variation in the spatial distribution of the neural response. To investigate this, we examined two complementary measures of spatial consistency between participants (**Jaccard Index**; **Voxel-wise Overlap**).

#### Preprocessing and Statistical Modeling

fMRI data were preprocessed using fMRIPrep v1.1.1 (Esteban et al., 2019), a pipeline developed by the Center for Reproducible Neuroscience.

Preprocessing steps included motion correction, correction for signal inhomogeneity, skull-stripping, segmentation, co-registration, and spatial normalization to the Montreal Neurological Institute (MNI)- 152 brain atlas. Preprocessed images were smoothed in SPM12 at 6mm FWHM. First level modeling was performed using SPM12. Individual regressors for each condition (INTEREST, NEUTRAL, BACKWARDS, and button press) were included in the model. Individual TRs were marked as outliers if they had greater than 1mm of framewise displacement. We included one regressor per outlier volume in the first level model, and we excluded participants with > 20% outlier volumes. The critical contrast (INTEREST > NEUTRAL) was created to examine regions showing greater activation for personally-interesting than neutral narratives.

#### Region of Interest Analyses

To determine whether personal interest activated language regions specifically, parameter estimates for each condition were extracted from *a priori* regions of interest (ROIs) known to be important for language processing (Fedorenko et al., 2010). These ROIs are based on an atlas comprised of functional data from 803 participants during language tasks and reflect regions wherein a high proportion of participants showed overlap in activation patterns (Lipkin et al., 2022). From this atlas, we selected eight parcels that contain canonical language regions: left IFGorb, left IFG, left MFG, left AntTemp, left PostTemp, left AngG, right cerebellum lobule VI, and right cerebellum Crus I/II (**see below**). Responses were averaged across all voxels within a given parcel. Linear mixed-effects models were run in R using the lme4 package. To determine if there was an effect of condition (INTEREST, NEUTRAL, BACKWARDS) across this pre-defined language network, we used:

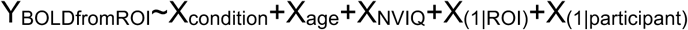

with participant and ROI as random factors to account for repeated measures, controlling for chronological age and nonverbal IQ (KBIT matrices). Post-hoc tests were then conducted using the lsmeans package in R.

#### Task-based Functional Connectivity Analysis

Task-based functional connectivity analyses were conducted using the CONN toolbox (Whitfield-Gabrieli & Nieto-Castanon, 2012) as implemented in MATLAB to determine whether personal interest altered the functional architecture of the language network. ROIs consisted of language network ROIs from (Lipkin et al., 2022), as used for the univariate analysis (i.e., left IFGorb, left IFG, left MFG, left AntTemp, left PostTemp, left AngG, right cerebellum lobule VI, and right cerebellum Crus I/II). FMRI data were preprocessed (realigned, normalized, spatially smoothed at 6mm FWHM), high-pass filtered (0.01Hz), and denoised using CompCor (Behzadi et al., 2007) and participant-specific motion parameters and outliers. First-level analyses were performed as a weighted-GLM. At the second level, ROI-ROI analyses were conducted to determine whether personal interest changed interactions between nodes of the language network as compared to neutral stories. Functional Network Connectivity multivariate statistics (Jafri et al., 2008) were used to analyze the set (network) of connections, thresholded at the cluster level with a FDR-corrected p<0.05 with a post-hoc uncorrected connection-level p<0.05.

Within-group matrices representing correlation values were visualized using the CONN toolbox.

#### Whole Brain fMRI Analyses

Whole-brain analyses were conducted to determine whether personal interests activated regions outside of canonical language regions. Group-level modeling was performed using SPM12. First-level maps from several contrasts of interest (NEUTRAL, INTEREST, INTEREST > NEUTRAL) were brought up to a second-level analyses wherein one-sample t-tests were used to determine regions for which activation in the condition of interest was greater than baseline. Group maps were thresholded at an uncorrected voxel p < 0.001, with a cluster correction for multiple comparisons (FWE < 0.05).

#### Jaccard Index

To measure the similarity in the spatial distribution of the whole brain responses between individual participants, we calculated the Jaccard Index (JI) (Jaccard, 1908) between each pair of participants for the INTEREST and NEUTRAL conditions. JI is a measure of the proportion of voxels in common between two conditions (the intersection of INTEREST and NEUTRAL contrast maps) with respect to the total of activation voxels identified by either condition (union of INTEREST and NEUTRAL contrast maps). In each condition, each participant’s first-level activation map was thresholded at t>2.3 and binarized, then JI was calculated across the whole brain using the formula: Mapi∩Mapj/Mapi∪Mapj for all possible pairs of maps *i* and *j*.

#### Voxel-wise Overlap

A group-constrained subject specific (GCSS) approach was used to assess consistency and spatial overlap in activation patterns across different conditions (Fedorenko et al., 2010; Scott & Perrachione, 2019). Each participant’s statistical parametric map of INTEREST > NEUTRAL was thresholded voxelwise at *p*<0.001 (uncorrected) and binarized. Binarized maps were overlaid to create a probability map of regions engaged more by INTEREST than NEUTRAL narratives, which were smoothed at 6mm FWHM and thresholded voxelwise at n=2 participants.

Probability maps reflect the number of participants showing overlap in a particular voxel.

#### Preregistration

The main hypotheses for the current paper were included as part of a broader preregistration in 2018 for a study investigating the neural correlates of personal interest in visual, reward, and language domains in neurotypical and autistic children: https://osf.io/nr3gk. Though beyond the scope of the current paper, the planned study included additional groups (e.g., a group of autistic children with strong interests and a group of neurotypical children without strong interests), as well as a video task and associated analyses that are not presented in the current paper (see **Supplementary Materials** for details on the videos task). For the analysis of the personal interest language task, we deviated from the preregistration by not using subject-specific functional ROIs (NEUTRAL>BACKWARDS), as we did not end up having sufficient data to independently define and test individually-defined regions for all participants. Instead, we used *a priori* ROIs and whole brain analyses. We did not test hypotheses related to associations between neural and behavioral measures due to smaller than anticipated sample sizes as a result of the COVID-19 pandemic and related data acquisition and personnel limitations.

## RESULTS

Our sample included 20 children, each with a strong specific interest. There was substantial variation among topics of interest between children, ranging from soccer to Minecraft to musicals (**Table 1**). We created a set of personalized narratives for each individual based on their interest (INTEREST) and presented these narratives in the scanner, interspersed with non-personalized (NEUTRAL) narratives about nature and time-reversed (BACKWARDS) narratives. The INTEREST and NEUTRAL narratives were closely matched on several linguistic and paralinguistic features (see Methods: Personalized Stimuli Creation and Supplementary Materials).

### Personally-interesting narratives increase activation in language regions

To determine whether personally-interesting narratives modulated activation in regions canonically associated with language processing, we extracted functional responses from each task condition (INTEREST, NEUTRAL, BACKWARDS) from eight *a priori* left frontal, temporal, parietal, and right cerebellar regions of interest (ROIs) that are consistently activated by language (Fedorenko et al., 2010). A mixed-effects model determined that brain activation differed by condition (F(2,451) = 121.43, p < 0.001). There were no effects of age or nonverbal IQ. As expected, comprehensible, generic narratives activated language regions more than incomprehensible, backwards narratives (NEUTRAL>BACKWARDS: t(451) = 4.89, p < 0.001). In addition, activation was higher for personally-interesting narratives than for both non-personalized (INTEREST>NEUTRAL: t(451) = 10.37, p < 0.001) and backwards (INTEREST>BACKWARDS: t(451) = 15.26, p < 0.001) narratives (**Figure 2A**).

**Figure 2:**
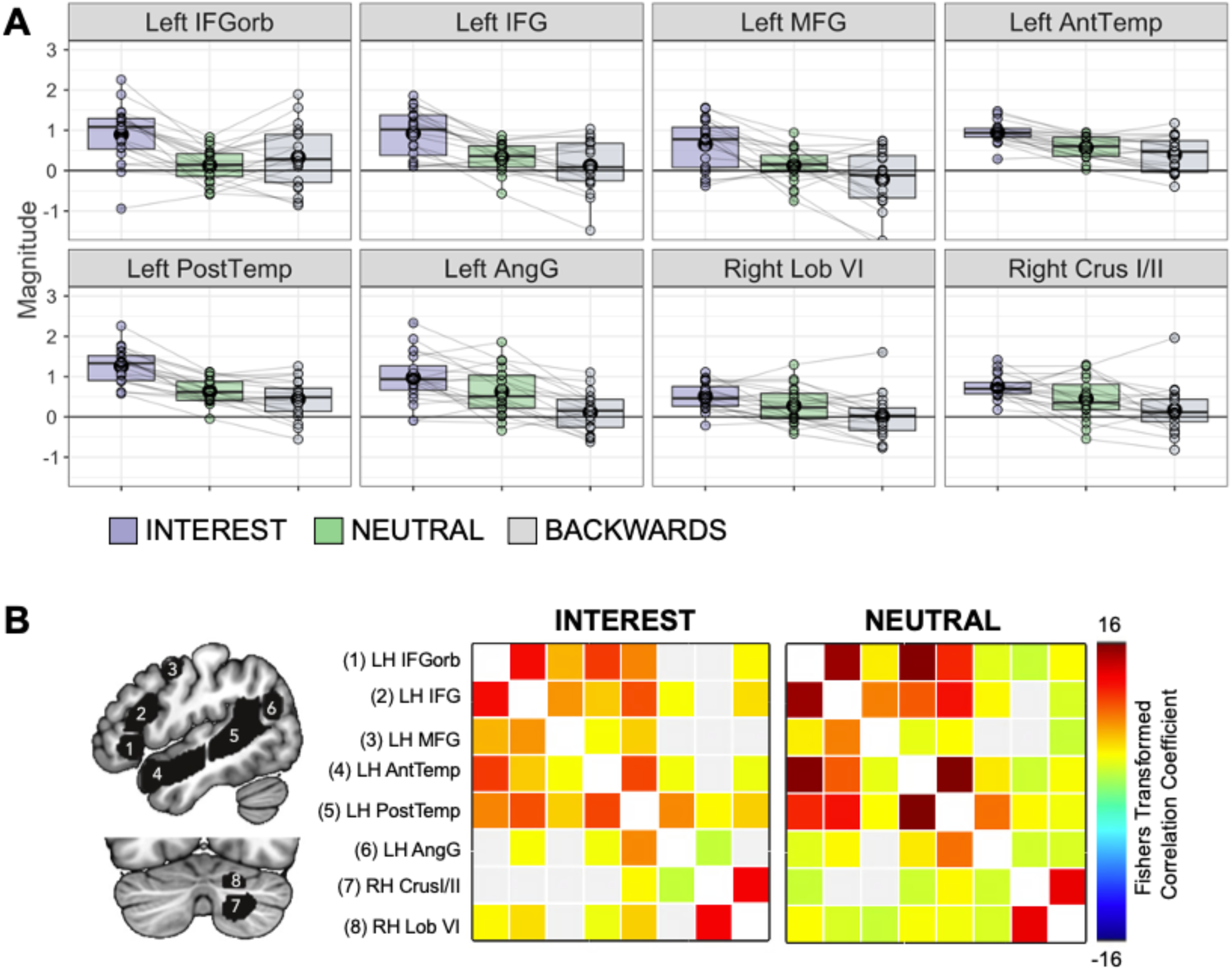
Response within a priori language regions of interest. (A) Boxplots show average magnitude per condition, per participant, in each a priori language region of interest (ROI). Circles represent individual participants; gray lines connect individual participants across conditions. Black circles represent the mean. (B) LEFT: A priori language regions of interest (Lipkin et al., 2022). RIGHT: Functional connectivity between language ROIs for the INTEREST and NEUTRAL conditions. Color bar represents Fisher-transformed bivariate correlation coefficients, thresholded using standard Functional Network Connectivity (FNC) parametric multivariate statistics with a FDR-corrected p<0.05 cluster-level threshold, together with a post-hoc uncorrected p<0.05 connection-level threshold.

We also assessed whether listening to personally-interesting narratives impacted functional connectivity between language regions. Inter-region correlations among canonical language regions did not differ significantly for the INTEREST compared to the NEUTRAL condition (**Figure 2B**).

Together, these results suggest that modulating interest impacts the magnitude of responses within language regions but does not impact how language regions interact with each other or alter the functional architecture of the language system.

Unlike generic language stimuli, personally-interesting narratives featured topics that were highly motivating and familiar for each child. We therefore conducted unbiased, whole-brain analyses to examine whether personally-interesting narratives elicited higher activation than neutral narratives outside canonical language regions as well. Whole-brain analyses confirmed that personally- interesting narratives increased activation in canonical language regions (e.g., bilateral superior and middle temporal gyri, bilateral inferior frontal gyrus, and right cerebellum) in both individual participants (**Figure 3A**; for individual responses in all participants see **Supplementary** Figure 1) and at the group level (**Figure 3B)**. Whole-brain analyses also revealed that personally-interesting narratives increased activation in regions associated with salience and reward (e.g., caudate, amygdala, nucleus accumbens, ventral tegmental area/substantia nigra) and self-referential processing (e.g., ventromedial and medial prefrontal cortex [vmPFC and mPFC], and precuneus/posterior cingulate) (**Figure 3B)**. Notably, though outside the scope of this paper, the INTEREST videos also elicited higher responses than the NEUTRAL videos in visual processing regions in the video task (see **Supplementary Materials** for group maps and discussion).

**Figure 3:**
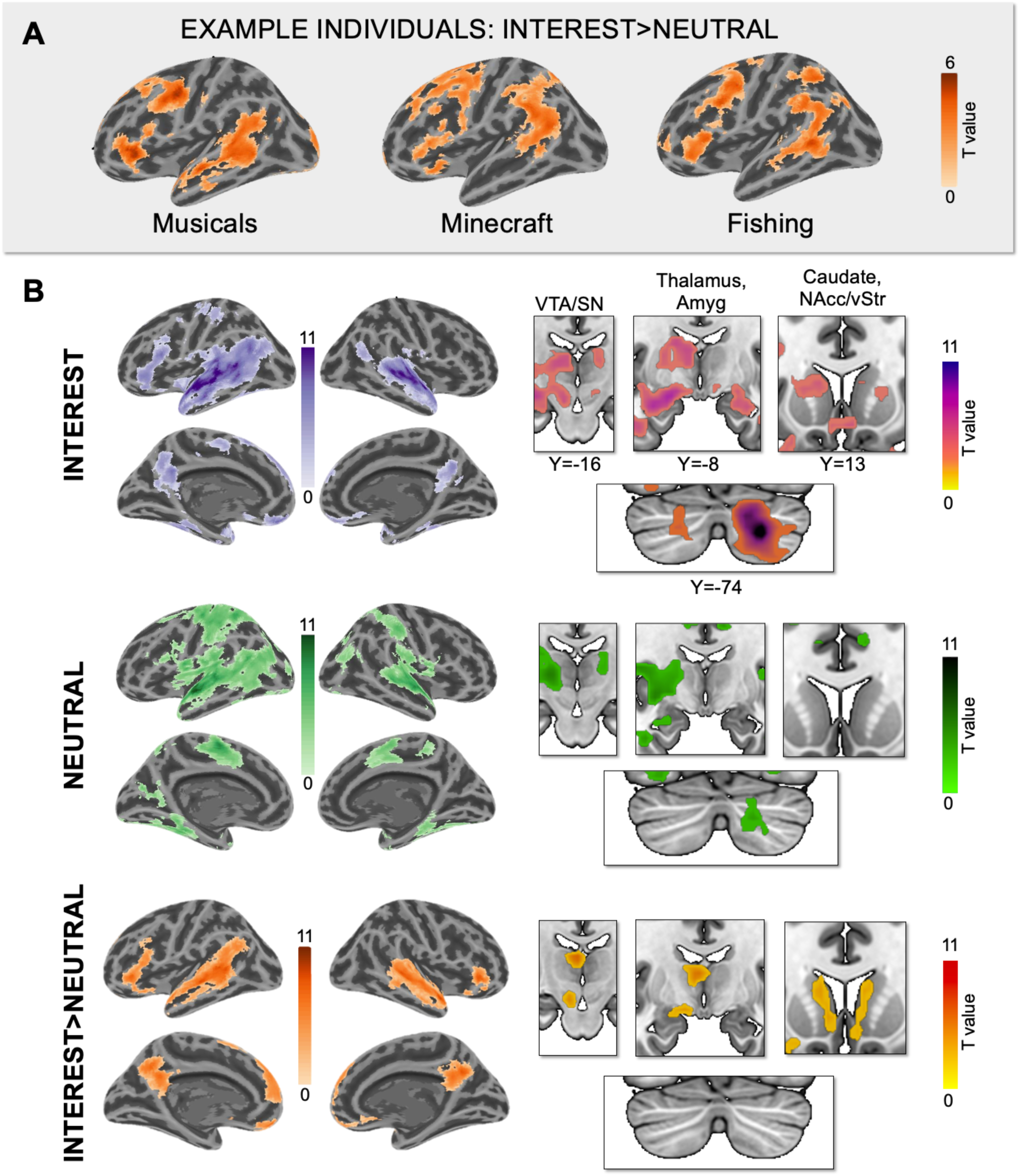
Personally-interesting narratives increase activation in the brain. (A) Individual activation maps from n=3 sample NT participants showing increased engagement of canonical language regions for personally interesting versus generic narratives (INTEREST > NEUTRAL single-subject maps thresholded at p<0.01, FWE cluster correction <0.05). (B) Whole-brain activation maps for the INTEREST, NEUTRAL, and INTEREST>NEUTRAL contrasts (threshold: p<0.001, FWE corrected at p<.05). Color bars reflect t-values.

### Personalized narratives activate brains more consistently despite idiosyncratic topics

A concern with personalization is that using different materials will give rise to discrepant patterns of activation across individuals. Intuitively, given that the non-personalized narratives were identical across individuals, one might expect more consistent patterns of activation across participants. To quantify the degree of spatial consistency in whole brain responses across individuals, we calculated the Jaccard Index (JI) (Jaccard, 1908) – a measure of spatial correspondence between two activation maps independent of activation magnitude – for the INTEREST and NEUTRAL conditions for each pair of subjects. The JI was significantly higher for the INTEREST condition than the NEUTRAL condition (t(19)=10.94, p<0.001; **Figure 4A**), indicating greater intersubject similarity in the spatial distribution of brain responses for personalized stimuli, despite the fact that they were idiosyncratic. We also found greater intersubject spatial consistency at the voxel level for the INTEREST compared to the NEUTRAL narratives (**Figure 4B**), with certain voxels *only* showing intersubject overlap in the INTEREST condition. These results suggest that even though the stimuli were idiosyncratic, ranging in topic from train lines to video games, activation patterns for personally-interesting narratives were *more consistent* in a greater number of voxels across participants than for neutral narratives.

**Figure 4:**
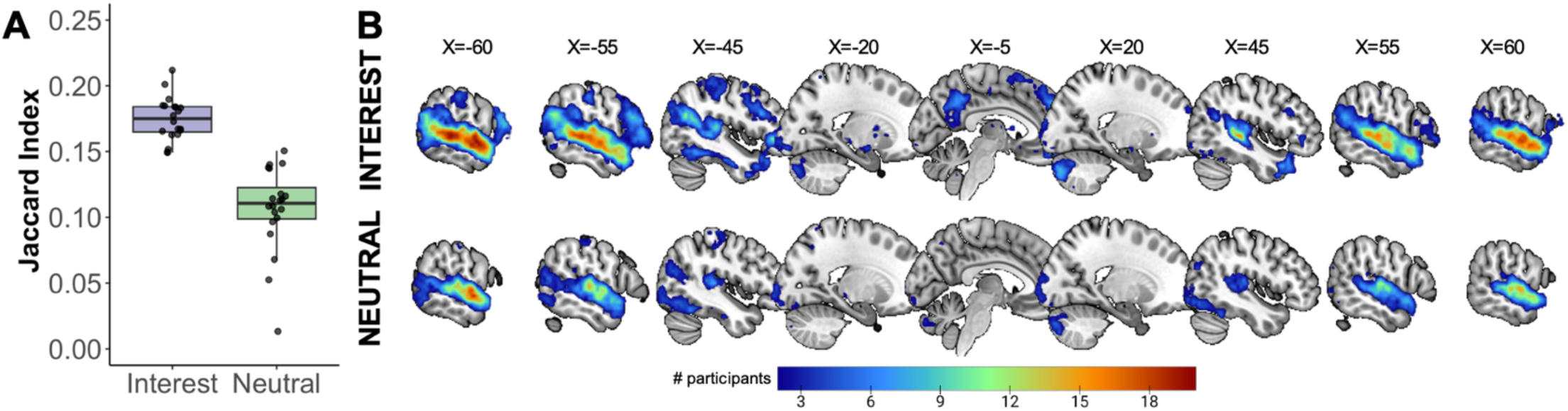
More consistent responses to personally-interesting than neutral language in neurotypical children. (A) Jaccard indices quantifying intersubject consistency (for each pair of participants) in whole brain activation patterns for the INTEREST (purple) and NEUTRAL (green) conditions. Each dot represents the mean JI for each participant. (B) Probability maps for INTEREST and NEUTRAL, showing the number of participants (out of 20 total) with overlapping activation.

## DISCUSSION

Some experiences are highly specific to the individual, making them difficult to study using traditional neuroimaging approaches that employ generic stimuli. Here, in 20 children, we took a novel approach to examine the effects of personal interests on language processing in the brain. To respect the idiosyncratic nature of interests, we created personalized neuroimaging experiments for each child using narratives written about their own interests.

### Activation in the brain is amplified for personally-interesting language

Personally-interesting narratives increased activation more than neutral narratives in several neocortical and cerebellar regions typically activated by language in both adults and children (Enge et al., 2020; Fedorenko et al., 2011; Friederici, 2011; Hiersche et al., 2024; Ozernov-Palchik et al., 2024; Price, 2010). This difference emerged at the level of individual participants, even though both conditions were high-level narratives and closely matched on low-level aspects of language.

Importantly, the functional architecture of the language network (functional connections between major hubs) was not affected by interest: interest modulated the magnitude of the response in language regions without changing how they communicated with each other. Prior studies have shown that some features of language stimuli (e.g., how surprising or predictable, semantically- plausible, grammatical, or challenging a sentence is) can upregulate or downregulate activation in language regions (e.g., D’Mello et al., 2020; Price, 2010; Shain et al., 2022; Tuckute et al., 2024; Willems et al., 2016), even though language regions are not sensitive to non-linguistic experimental manipulations (e.g., Fedorenko et al., 2011, 2020). Although “interest” is not typically considered a linguistic feature, our results show that interests do influence the brain’s response in regions associated with language processing. Notably, interest – unlike the other linguistic features mentioned above – is *determined by the individual* and is not intrinsic to the stimuli, and thus cannot be accounted for without involving participants in study design.

We also found that personally-interesting narratives elicited higher activation than generic narratives in areas associated with reward and self-reference. Although not considered “language” regions (i.e., not specifically and selectively involved in language processing), several of these regions are recruited by typical language processing tasks (Lipkin et al., 2022). It is possible that the personally-interesting narratives brought online regions that neuroimaging paradigms using generic language stimuli only detect with very large sample sizes (e.g., Lipkin et al., 2022) or regions that may be under-reported in studies of language (e.g., Janacsek et al., 2022). Another possibility is that the observed activation in areas such as the amygdala, ventral tegmental area, and ventral striatum/caudate reflect the highly rewarding and salient nature of topics of interest.

Canonical default-mode network (DMN) regions such as mPFC and precuneus/posterior cingulate were also more active for the personally-interesting language stimuli, perhaps due to the inherently self-referential aspects of interest (e.g., interests are closely linked to the self – both how individuals view themselves and the way they project to the world (Harackiewicz et al., 2016; Renninger & Hidi, 2015)). Importantly, engagement of reward and self-referential regions has been noted in previous studies that examined the neural basis of personal relevance (Abraham, 2013; Abraham & Cramon, 2009) and that experimentally manipulated personal relevance in linguistic stimuli (e.g., participants’ mothers’ voices versus unfamiliar voices (Abrams et al., 2016, 2022)).

Activation of the DMN and basal ganglia are also consistently noted in language studies that use coherent narratives, similar to the narratives used in the current study, rather than isolated sentences (Lerner et al., 2011; Silbert et al., 2014; Zadbood et al., 2017). Based on this prior literature, it is possible that increased activation in these regions is related to one or more unique aspects of the personally-interesting narratives (e.g., content, greater personal relevance, salience, reward).

A key question emerges from these results: *why* does the brain respond more to the personally-interesting narratives? Our results are in line with several related studies that find that factors that increase engagement and attention to language (Cohen et al., 2021; Grall et al., 2021; Song et al., 2021) – and even individually-varying factors like personality traits (Finn et al., 2018) – modulate activity in the brain regions such as the ones we identified in the current study. However, unlike these other studies which used uniform stimuli but took advantage of naturally-occuring individual variation, the current study used *personalized* stimuli to match individuals on a factor which typically varies – interest. Critically, by also closely matching our experimental conditions on several features (e.g., speaker, emotional valence, and parts of speech), this study design enabled us to isolate the specific effects of interest as the one condition we experimentally manipulated. Of course, to do this, we had to cede some control over low-level linguistic features in some participants (e.g., using proper nouns or low-frequency words). In our case, the trade-off between personalization and perfect control over linguistic variables is analogous to the trade-offs made in other studies that prioritize control over intrinsic features of the stimuli but may overlook the impact of personal experience and interest.

### Personalizing narratives to the individual reveals unexpected benefits

Across the field of neuroimaging, there has been a growing awareness that considering only group-level effects limits the validity and translational potential of results, prompting a movement toward studying individual differences. To this end, researchers have predominantly relied on ‘precision’ neuroimaging, which often involves collecting large amounts of data on a small number of individuals to reliably characterize individual differences in brain responses to the same stimuli (Gabrieli et al., 2015; Gordon et al., 2017; Gratton et al., 2020, 2022; Kraus et al., 2023).

However, results from the current study suggest that an alternative approach may be to incorporate individual differences at the level of stimuli design. Critically, instead of leading to more heterogeneous patterns of activation across individuals, personalized narratives actually elicited *more consistent* patterns of brain activation across individuals than generic narratives. This result suggests that while tight experimental control mitigates low-level differences across participants, it could introduce confounds at other levels which could both limit our power to detect brain responses, and affect our interpretation of brain differences. Furthermore, with increasing awareness of individual variablity in the precise location of functional regions in the human brain (Fedorenko et al., 2010; Saxe et al., 2006), our results suggest that there should be a similar acknowledgement of how individual differences in participants’ *subjective experiences* of experimental stimuli affect interpretation of neuroimaging results. Indeed, prior studies have found that researcher labels of stimuli do not always capture participants’ subjective experiences of those stimuli, and that using participants’ own reports rather than experimenter labels can explain more variance in brain activity (Varrier & Finn, 2022). This issue has been indirectly addressed in a handful of neuroimaging studies that have successfully used personalized stimuli when studying phenomena that are more clearly personal such as food cravings, memories, or content dimensions of spontaneous thoughts (Bainbridge & Baker, 2022; Kim et al., 2024; Tomova et al., 2020). We propose that individual-level factors are an important consideration when evaluating brain responses across various cognitive domains.

Personalized stimuli may be particularly beneficial for certain populations, such as young children and individuals with neurodevelopmental delays or psychiatric conditions, who have historically been challenging to study using fMRI. Utilizing interest to drive attention has been part of educational philosophy for at least a century – i.e., if children are interested in a topic, they will engage with it more meaningfully and be motivated to try hard things (Krapp et al., 1992; Renninger et al., 2014; Renninger & Hidi, 2015; Walkington & Bernacki, 2014) – and this phenomenon has been both underutilized and underexplored in neuroimaging. Scaffolding communicative interactions and interventions around topics of personal interest has also been shown to support social and communication skills for autistic children (Arunachalam et al., 2024; Baker et al., 1998; Boyd et al., 2007; Charlop-Christy & Haymes, 1998; Harrop et al., 2019; Lizon et al., 2023; Suskind, 2014), and even to modulate brain activation (Cascio et al., 2014; Dichter et al., 2012; Foss-Feig et al., 2016; Kohls et al., 2018; Pierce & Redcay, 2008). Finally, using more naturalistic stimuli has been shown to improve data quality in other pediatric samples (Cantlon, 2020; Redcay & Moraczewski, 2020; Richardson et al., 2018; Vanderwal et al., 2019), and our results further support this approach. In particular, interest may be an additional dimension that could increase engagement and more accurately capture the organization and function of the brain.

### Limitations and Future Directions

This study was limited in a few ways that future research might aim to address. Because the interest narratives were personalized to individuals, the neutral and interest conditions were not perfectly matched on all linguistic dimensions (despite being matched on several low-level and lexical features). Further, the content of the narratives was variable – some narratives had social content, some involved more physical actions, some were fictional, etc. Although these factors can modulate brain activation, they are unlikely to drive the results we observed as the contrast of interest was robust at the level of individual participants (including those for whom differences in linguistic properties between the interest and neutral narratives were minimal, and across a wide variety of topics). It is also difficult to determine whether our results are a product of *personalization* or general increased interest in the stimuli. It is worth noting that the nature narratives were not *un-*interesting to participants, and interest is a continuous dynamic variable. Finally, the experimental design was passive, the critical language fMRI task followed exposure to videos relating to each child’s interest, and the neutral narratives were interspersed with highly engaging personalized narratives. These factors could all have modulated attention (e.g., children could have attended more to the INTEREST condition because the topic was more engaging, or alternatively attended more to the NEUTRAL condition because content of the narratives was about a topic they did not think about as often). However, both narrative conditions were designed to be engaging, and indeed, both conditions robustly activated the brain.

There are multiple followup directions to this research that would provide insight into the robust condition differences we observed. For example, future studies could employ a more active task design to more explicitly match level of engagement across conditions. Additionally, while in this paper we focused on spatial consistency across participants, another exciting future direction is to examine the effects of interest on temporal consistency of neural responses using intersubject correlation (ISC) analyses (Hasson et al., 2004). Finally, future studies should further manipulate both linguistic and individual-level (i.e., interest) features of linguistic stimuli in different populations to more precisely understand their respective contributions to functional language responses in the brain.

Future studies should also examine the effects of interest on other cognitive domains.

### Conclusions

In sum, this study highlights that personal interests modulate language function in the brains of children. For the first time, these results also illustrate that personalizing stimuli is a feasible way to evoke consistent brain responses. Finally, this study underscores the notion that the language system is sensitive to the content of narratives and that personal interests might be a powerful tool for probing the scope and functionality of brain networks.

## Data and Code Availability

Data and materials are available on OSF (https://osf.io/dh3wq/).

## Author Contributions

AMD and JDEG conceptualized the study. AMD, HAO, and KTJ designed the experiment. HAO recorded all audio stimuli. HAO, AMD, KTJ, IRF, and SN collected data. HAO and AMD analyzed the data. HAO and AMD wrote the initial draft of the manuscript. KTJ and JDEG edited the manuscript.

## Funding

This research was supported by the Hock E. Tan and K. Lisa Yang Center for Autism Research at MIT, Seth Klarman and Paul Gannon (to JDEG), NIH F32 MH117933 and Simons Center for the Social Brain Postdoctoral Fellowship (to AMD), NSF Graduate Research Fellowship Program #1745302 (to HAO), and MIT Hugh Hampton Young Memorial Fellowship and MIT Media Lab Learning Innovation Fellowship (to KTJ).

## Declaration of Competing Interests

The authors declare no competing financial interests.

## Acknowledgments

We thank Cindy Li for assistance with recruiting, coordinating, and testing with our autism group. We also thank Hannah Grotzinger for task programming assistance, Caitlin Malloy for ADOS support, and our undergraduate and high school research assistants who assisted with various aspects of the project, including Jimmy Chen, Nicole Dundas, Insha Merchant, Rucha Kelkar, Alana Kalehua, and Hillary Jean-Gilles. We are grateful to Atsushi Takahashi and Steve Shannon from the Athinoula A. Martinos Imaging Center at MIT. We thank the participants and families for making this research possible. Finally, we thank Ron Suskind, whose experience with his son Owen inspired this research.

## Notes

https://osf.io/dh3wq/

### Competing Interest Statement

The authors have declared no competing interest.

### Summary of Updates

The manuscript has been revised to add additional analyses, clarify the argument, and update the focus of the paper.

## REFERENCES

Abraham, A. (2013). The World According to Me: Personal Relevance and the Medial Prefrontal Cortex. Frontiers in Human Neuroscience, 7. https://www.frontiersin.org/articles/10.3389/fnhum.2013.00341

Abraham, A., & Cramon, D. Y. von. (2009). Reality = Relevance? Insights from Spontaneous Modulations of the Brain’s Default Network when Telling Apart Reality from Fiction. PLOS ONE, 4(3), e4741. 10.1371/journal.pone.0004741

Abrams, D. A., Chen, T., Odriozola, P., Cheng, K. M., Baker, A. E., Padmanabhan, A., Ryali, S., Kochalka, J., Feinstein, C., & Menon, V. (2016). Neural circuits underlying mother’s voice perception predict social communication abilities in children. Proceedings of the National Academy of Sciences, 113(22), 6295–6300. 10.1073/pnas.1602948113

Abrams, D. A., Mistry, P. K., Baker, A. E., Padmanabhan, A., & Menon, V. (2022). A Neurodevelopmental Shift in Reward Circuitry from Mother’s to Nonfamilial Voices in Adolescence. Journal of Neuroscience, 42(20), 4164–4173. 10.1523/JNEUROSCI.2018-21.2022

Arunachalam, S., Steele, A., Pelletier, T., & Luyster, R. (2024). Do focused interests support word learning? A study with autistic and nonautistic children. Autism Research, n/a(n/a). 10.1002/aur.3121

Bainbridge, W. A., & Baker, C. I. (2022). Multidimensional memory topography in the medial parietal cortex identified from neuroimaging of thousands of daily memory videos. Nature Communications, 13(1), Article 1. 10.1038/s41467-022-34075-1

Baker, M. J., Koegel, R. L., & Koegel, L. K. (1998). Increasing the Social Behavior of Young Children with Autism Using Their Obsessive Behaviors. Journal of the Association for Persons with Severe Handicaps, 23(4), 300–308. 10.2511/rpsd.23.4.300

Behzadi, Y., Restom, K., Liau, J., & Liu, T. T. (2007). A Component Based Noise Correction Method (CompCor) for BOLD and Perfusion Based fMRI. NeuroImage, 37(1), 90–101. 10.1016/j.neuroimage.2007.04.042

Boyd, B. A., Conroy, M. A., Mancil, G. R., Nakao, T., & Alter, P. J. (2007). Effects of Circumscribed Interests on the Social Behaviors of Children with Autism Spectrum Disorders. Journal of Autism and Developmental Disorders, 37(8), 1550–1561. 10.1007/s10803-006-0286-8

Cantlon, J. F. (2020). The balance of rigor and reality in developmental neuroscience. NeuroImage, 216, 116464. 10.1016/j.neuroimage.2019.116464

Cascio, C. J., Foss-Feig, J. H., Heacock, J., Schauder, K. B., Loring, W. A., Rogers, B. P., Pryweller, J. R., Newsom, C. R., Cockhren, J., Cao, A., & Bolton, S. (2014). Affective neural response to restricted interests in autism spectrum disorders. Journal of Child Psychology and Psychiatry, 55(2), 162–171. 10.1111/jcpp.12147

Charlop-Christy, M. H., & Haymes, L. K. (1998). Using Objects of Obsession as Token Reinforcers for Children with Autism. Journal of Autism and Developmental Disorders, 28(3), 189–198. 10.1023/A:1026061220171

Cohen, L., Salondy, P., Pallier, C., & Dehaene, S. (2021). How does inattention affect written and spoken language processing? Cortex, 138, 212–227. 10.1016/j.cortex.2021.02.007

Dichter, G. S., Felder, J. N., Green, S. R., Rittenberg, A. M., Sasson, N. J., & Bodfish, J. W. (2012). Reward circuitry function in autism spectrum disorders. Social Cognitive and Affective Neuroscience, 7(2), 160–172. 10.1093/scan/nsq095

D’Mello, A. M., Centanni, T. M., Gabrieli, J. D. E., & Christodoulou, J. A. (2020). Cerebellar Contributions to Rapid Semantic Processing in Reading. Brain and Language, 208, 104828. 10.1016/j.bandl.2020.104828

Enge, A., Friederici, A. D., & Skeide, M. A. (2020). A meta-analysis of fMRI studies of language comprehension in children. NeuroImage, 215, 116858. 10.1016/j.neuroimage.2020.116858

Esteban, O., Markiewicz, C. J., Blair, R. W., Moodie, C. A., Isik, A. I., Erramuzpe, A., Kent, J. D., Goncalves, M., DuPre, E., Snyder, M., Oya, H., Ghosh, S. S., Wright, J., Durnez, J., Poldrack, R. A., & Gorgolewski, K. J. (2019). fMRIPrep: A robust preprocessing pipeline for functional MRI. Nature Methods, 16(1), Article 1. 10.1038/s41592-018-0235-4

Fedorenko, E., Behr, M. K., & Kanwisher, N. (2011). Functional specificity for high-level linguistic processing in the human brain. Proceedings of the National Academy of Sciences, 108(39), 16428–16433. 10.1073/pnas.1112937108

Fedorenko, E., Blank, I. A., Siegelman, M., & Mineroff, Z. (2020). Lack of selectivity for syntax relative to word meanings throughout the language network. Cognition, 203, 104348. 10.1016/j.cognition.2020.104348

Fedorenko, E., Hsieh, P.-J., Nieto-Castañón, A., Whitfield-Gabrieli, S., & Kanwisher, N. (2010). New Method for fMRI Investigations of Language: Defining ROIs Functionally in Individual Subjects. Journal of Neurophysiology, 104(2), 1177–1194. 10.1152/jn.00032.2010

Fink, R. P. (1995). Successful Dyslexics: A Constructivist Study of Passionate Interest Reading. Journal of Adolescent & Adult Literacy, 39(4), 268–280.

Finn, E. S., Corlett, P. R., Chen, G., Bandettini, P. A., & Constable, R. T. (2018). Trait paranoia shapes inter-subject synchrony in brain activity during an ambiguous social narrative. Nature Communications, 9(1), Article 1. 10.1038/s41467-018-04387-2

Foss-Feig, J. H., McGugin, R. W., Gauthier, I., Mash, L. E., Ventola, P., & Cascio, C. J. (2016). A functional neuroimaging study of fusiform response to restricted interests in children and adolescents with autism spectrum disorder. Journal of Neurodevelopmental Disorders, 8(1). 10.1186/s11689-016-9149-6

Friederici, A. D. (2011). The Brain Basis of Language Processing: From Structure to Function. Physiological Reviews, 91(4), 1357–1392. 10.1152/physrev.00006.2011

Gabrieli, J. D. E., Ghosh, S. S., & Whitfield-Gabrieli, S. (2015). Prediction as a Humanitarian and Pragmatic Contribution from Human Cognitive Neuroscience. Neuron, 85(1), 11–26. 10.1016/j.neuron.2014.10.047

Gordon, E. M., Laumann, T. O., Gilmore, A. W., Newbold, D. J., Greene, D. J., Berg, J. J., Ortega, M., Hoyt-Drazen, C., Gratton, C., Sun, H., Hampton, J. M., Coalson, R. S., Nguyen, A. L., McDermott, K. B., Shimony, J. S., Snyder, A. Z., Schlaggar, B. L., Petersen, S. E., Nelson, S. M., & Dosenbach, N. U. F. (2017). Precision Functional Mapping of Individual Human Brains. Neuron, 95(4), 791–807.e7. 10.1016/j.neuron.2017.07.011

Grall, C., Tamborini, R., Weber, R., & Schmälzle, R. (2021). Stories Collectively Engage Listeners’ Brains: Enhanced Intersubject Correlations during Reception of Personal Narratives. Journal of Communication, 71(2), 332–355. 10.1093/joc/jqab004

Gratton, C., Kraus, B. T., Greene, D. J., Gordon, E. M., Laumann, T. O., Nelson, S. M., Dosenbach, N. U. F., & Petersen, S. E. (2020). Defining Individual-Specific Functional Neuroanatomy for Precision Psychiatry. Biological Psychiatry, 88(1), 28–39. 10.1016/j.biopsych.2019.10.026

Gratton, C., Nelson, S. M., & Gordon, E. M. (2022). Brain-behavior correlations: Two paths toward reliability. Neuron, 110(9), 1446–1449. 10.1016/j.neuron.2022.04.018

Harackiewicz, J. M., Smith, J. L., & Priniski, S. J. (2016). Interest Matters: The Importance of Promoting Interest in Education. Policy Insights from the Behavioral and Brain Sciences, 3(2), 220–227. 10.1177/2372732216655542

Harrop, C., Amsbary, J., Towner-Wright, S., Reichow, B., & Boyd, B. A. (2019). That’s what I like: The use of circumscribed interests within interventions for individuals with autism spectrum disorder. A systematic review. Research in Autism Spectrum Disorders, 57, 63–86. 10.1016/j.rasd.2018.09.008

Hasson, U., Nir, Y., Levy, I., Fuhrmann, G., & Malach, R. (2004). Intersubject Synchronization of Cortical Activity During Natural Vision. Science, 303(5664), 1634–1640. 10.1126/science.1089506

Hiersche, K. J., Schettini, E., Li, J., & Saygin, Z. M. (2024). Functional dissociation of the language network and other cognition in early childhood. Human Brain Mapping, 45(9), e26757. 10.1002/hbm.26757

Jaccard, P. (1908). Nouvelles recherches sur la distribution florale. Bull. Soc. Vaud. Sci. Nat., 44, 223–270.

Jafri, M. J., Pearlson, G. D., Stevens, M., & Calhoun, V. D. (2008). A method for functional network connectivity among spatially independent resting-state components in schizophrenia. NeuroImage, 39(4), 1666–1681. 10.1016/j.neuroimage.2007.11.001

Janacsek, K., Evans, T. M., Kiss, M., Shah, L., Blumenfeld, H., & Ullman, M. T. (2022). Subcortical Cognition: The Fruit Below the Rind. Annual Review of Neuroscience, 45(Volume 45, 2022), 361–386. 10.1146/annurev-neuro-110920-013544

Kaufman, A. S., & Kaufman, N. L. (2004). Kaufman Brief Intelligence Test (2nd ed.). American Guidance Service.

Kim, H. J., Lux, B. K., Lee, E., Finn, E. S., & Woo, C.-W. (2024). Brain decoding of spontaneous thought: Predictive modeling of self-relevance and valence using personal narratives. Proceedings of the National Academy of Sciences, 121(14), e2401959121. 10.1073/pnas.2401959121

Kohls, G., Antezana, L., Mosner, M. G., Schultz, R. T., & Yerys, B. E. (2018). Altered reward system reactivity for personalized circumscribed interests in autism. Molecular Autism, 9(1). 10.1186/s13229-018-0195-7

Krapp, A., Hidi, S., & Renninger, K. A. (1992). Interest, learning, and development. In K. A. Renninger, S. Hidi, & A. Krapp (Eds.), The role of interest in learning and development (pp. 3– 25). Lawrence Erlbaum Associates, Inc.

Kraus, B., Zinbarg, R., Braga, R. M., Nusslock, R., Mittal, V. A., & Gratton, C. (2023). Insights from personalized models of brain and behavior for identifying biomarkers in psychiatry. Neuroscience & Biobehavioral Reviews, 152, 105259. 10.1016/j.neubiorev.2023.105259

Lerner, Y., Honey, C. J., Silbert, L. J., & Hasson, U. (2011). Topographic Mapping of a Hierarchy of Temporal Receptive Windows Using a Narrated Story. Journal of Neuroscience, 31(8), 2906– 2915. 10.1523/JNEUROSCI.3684-10.2011

Lipkin, B., Tuckute, G., Affourtit, J., Small, H., Mineroff, Z., Kean, H., Jouravlev, O., Rakocevic, L., Pritchett, B., Siegelman, M., Hoeflin, C., Pongos, A., Blank, I. A., Struhl, M. K., Ivanova, A., Shannon, S., Sathe, A., Hoffmann, M., Nieto-Castañón, A., & Fedorenko, E. (2022). Probabilistic atlas for the language network based on precision fMRI data from >800 individuals. Scientific Data, 9(1), Article 1. 10.1038/s41597-022-01645-3

Lizon, M., Taels, L., & Vanheule, S. (2023). Specific interests as a social boundary and bridge: A qualitative interview study with autistic individuals. Autism, 13623613231193532. 10.1177/13623613231193532

Ozernov-Palchik, O., O’Brien, A. M., Lee, E. J., Richardson, H., Romeo, R., Lipkin, B., Small, H., Capella, J., Nieto-Castanon, A., Saxe, R., Gabrieli, J. D. E., & Fedorenko, E. (2024). Precision fMRI reveals that the language network exhibits adult-like left-hemispheric lateralization by 4 years of age (p. 2024.05.15.594172). bioRxiv. 10.1101/2024.05.15.594172

Pierce, K., & Redcay, E. (2008). Fusiform Function in Children with an Autism Spectrum Disorder Is a Matter of “Who.” Biological Psychiatry, 64(7), 552–560. 10.1016/j.biopsych.2008.05.013

Price, C. J. (2010). The anatomy of language: A review of 100 fMRI studies published in 2009. Annals of the New York Academy of Sciences, 1191(1), 62–88. 10.1111/j.1749-6632.2010.05444.x

Reber, R., Canning, E. A., & Harackiewicz, J. M. (2018). Personalized Education to Increase Interest. Current Directions in Psychological Science, 27(6), 449–454. 10.1177/0963721418793140

Recht, D. R., & Leslie, L. (1988). Effect of prior knowledge on good and poor readers’ memory of text. Journal of Educational Psychology, 80, 16–20. 10.1037/0022-0663.80.1.16

Redcay, E., & Moraczewski, D. (2020). Social cognition in context: A naturalistic imaging approach. NeuroImage, 216, 116392. 10.1016/j.neuroimage.2019.116392

Renninger, K. A., & Hidi, S. (2015). The Power of Interest for Motivation and Engagement. Routledge.

Renninger, K. A., Hidi, S., Krapp, A., & Renninger, A. (Eds.). (2014). The Role of interest in Learning and Development. Psychology Press. 10.4324/9781315807430

Richardson, H., Lisandrelli, G., Riobueno-Naylor, A., & Saxe, R. (2018). Development of the social brain from age three to twelve years. Nature Communications, 9(1), Article 1. 10.1038/s41467-018-03399-2

Saxe, R., Brett, M., & Kanwisher, N. (2006). Divide and conquer: A defense of functional localizers. NeuroImage, 30(4), 1088–1096. 10.1016/j.neuroimage.2005.12.062

Scott, T. L., & Perrachione, T. K. (2019). Common cortical architectures for phonological working memory identified in individual brains. NeuroImage, 202, 116096. 10.1016/j.neuroimage.2019.116096

Shain, C., Blank, I. A., Fedorenko, E., Gibson, E., & Schuler, W. (2022). Robust Effects of Working Memory Demand during Naturalistic Language Comprehension in Language-Selective Cortex. Journal of Neuroscience, 42(39), 7412–7430. 10.1523/JNEUROSCI.1894-21.2022

Shnayer, S. W. (1968). Some Relationships between Reading Interest and Reading Comprehension. https://eric.ed.gov/?id=ED022633

Silbert, L. J., Honey, C. J., Simony, E., Poeppel, D., & Hasson, U. (2014). Coupled neural systems underlie the production and comprehension of naturalistic narrative speech. Proceedings of the National Academy of Sciences, 111(43), E4687–E4696. 10.1073/pnas.1323812111

Song, H., Finn, E. S., & Rosenberg, M. D. (2021). Neural signatures of attentional engagement during narratives and its consequences for event memory. Proceedings of the National Academy of Sciences, 118(33), e2021905118. 10.1073/pnas.2021905118

Sparks, E., Schinkel, M. G., & Moore, C. (2017). Affiliation affects generosity in young children: The roles of minimal group membership and shared interests. Journal of Experimental Child Psychology, 159, 242–262. 10.1016/j.jecp.2017.02.007

Suskind, R. (2014). Life, Animated: A Story of Sidekicks, Heroes, and Autism.

Kingswell. Tomova, L., Wang, K. L., Thompson, T., Matthews, G. A., Takahashi, A., Tye, K. M., & Saxe, R. (2020). Acute social isolation evokes midbrain craving responses similar to hunger. Nature Neuroscience, 23(12), Article 12. 10.1038/s41593-020-00742-z

Tuckute, G., Sathe, A., Srikant, S., Taliaferro, M., Wang, M., Schrimpf, M., Kay, K., & Fedorenko, E. (2024). Driving and suppressing the human language network using large language models. Nature Human Behaviour, 1–18. 10.1038/s41562-023-01783-7

Vanderwal, T., Eilbott, J., & Castellanos, F. X. (2019). Movies in the magnet: Naturalistic paradigms in developmental functional neuroimaging. Developmental Cognitive Neuroscience, 36, 100600. 10.1016/j.dcn.2018.10.004

Varrier, R. S., & Finn, E. S. (2022). Seeing Social: A Neural Signature for Conscious Perception of Social Interactions. The Journal of Neuroscience, 42(49), 9211–9226. 10.1523/JNEUROSCI.0859-22.2022

Walkington, C., & Bernacki, M. L. (2014). Motivating Students by “Personalizing” Learning around Individual Interests: A Consideration of Theory, Design, and Implementation Issues. In Motivational Interventions (Vol. 18, pp. 139–176). Emerald Group Publishing Limited. 10.1108/S0749-742320140000018004

Whitfield-Gabrieli, S., & Nieto-Castanon, A. (2012). Conn: A Functional Connectivity Toolbox for Correlated and Anticorrelated Brain Networks. Brain Connectivity, 2(3), 125–141. 10.1089/brain.2012.0073

Willems, R. M., Frank, S. L., Nijhof, A. D., Hagoort, P., & van den Bosch, A. (2016). Prediction During Natural Language Comprehension. Cerebral Cortex, 26(6), 2506–2516. 10.1093/cercor/bhv075

Zadbood, A., Chen, J., Leong, Y. C., Norman, K. A., & Hasson, U. (2017). How We Transmit Memories to Other Brains: Constructing Shared Neural Representations Via Communication. Cerebral Cortex, 27(10), 4988–5000. 10.1093/cercor/bhx202

