## Supplemental Information for "Personalized Neuroimaging Reveals the Impact of Children’s Interests on Language Processing in the Brain"

**Supplementary Materials**

**PERSONALIZED LANGUAGE TASK**

***Supplementary Figure 1: Higher responses to personally interesting narratives in individual participants***

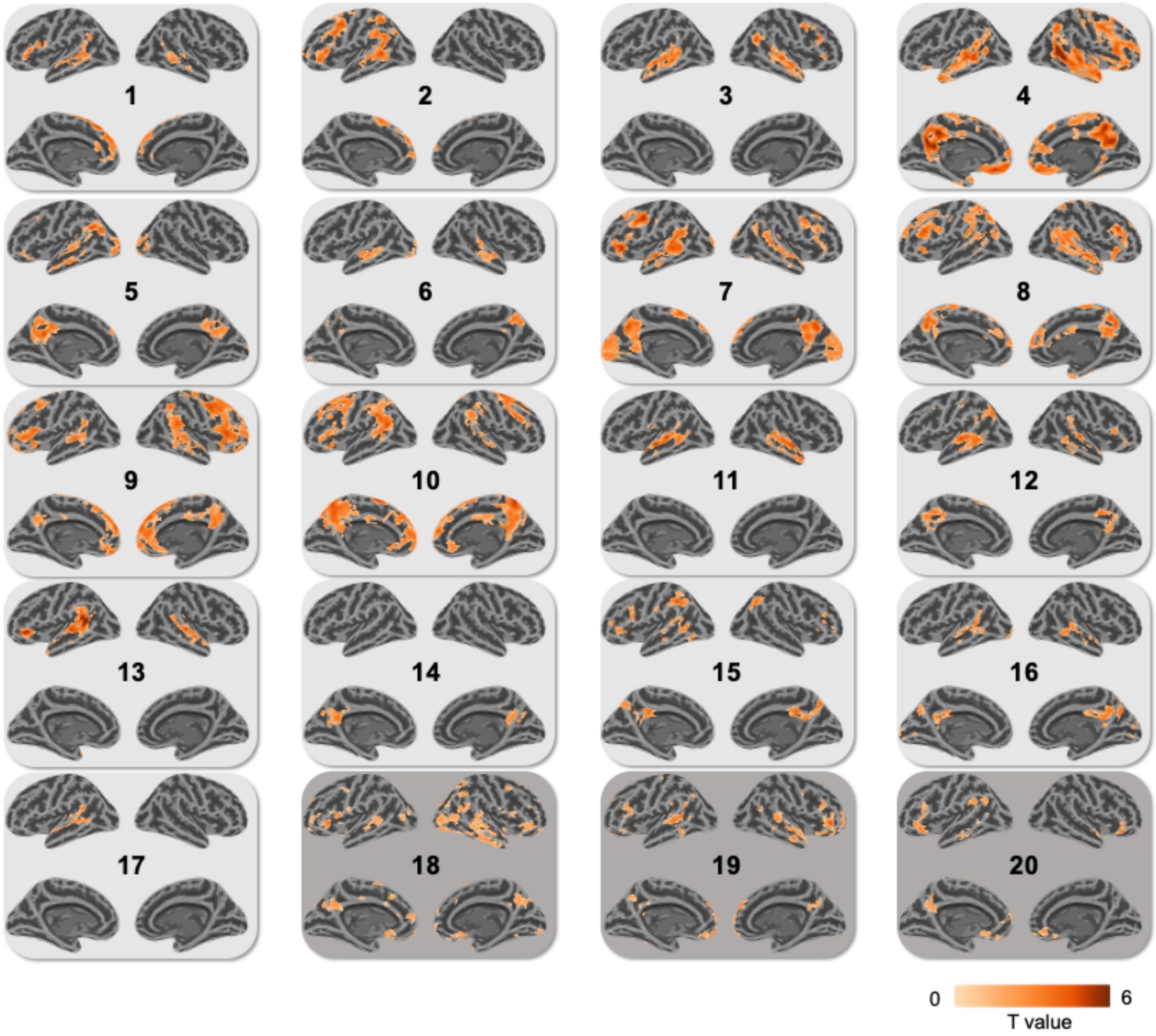


*Individual whole-brain responses from 20 participants to INTEREST>NEUTRAL language. N=17 participants visualized at p<0.01, FWE cluster p<0.05. N=3 participants did not show activation at this threshold or had activation that was not visible when projected onto surface space and are therefore visualized at p<0.05 uncorrected (dark grey boxes).*

**Stimuli characterization.**
Non-parametric pairwise comparisons were conducted using Wilcoxon rank-sum tests to identify specific differences between individual subjects' narratives and the neutral narratives. After applying a Bonferroni correction for multiple comparisons (adjusted $p<0.0025$ for 20 comparisons), no significant differences were found for the number of sentences, syllables per sentence, emotional valence, adverb usage, verb usage, and adjective usage across narratives. A few subjects showed significant differences for words per sentence, word count, and word frequency. While noun and proper noun usage were significantly different for several participants when examined separately, these differences were not observed when nouns and proper nouns were combined. Therefore, it is unlikely that the linguistic and paralinguistic features of the narratives were a driving factor in the overall results.

***Supplementary Figure 2: Linguistic and paralinguistic features for individual participants' stimuli***

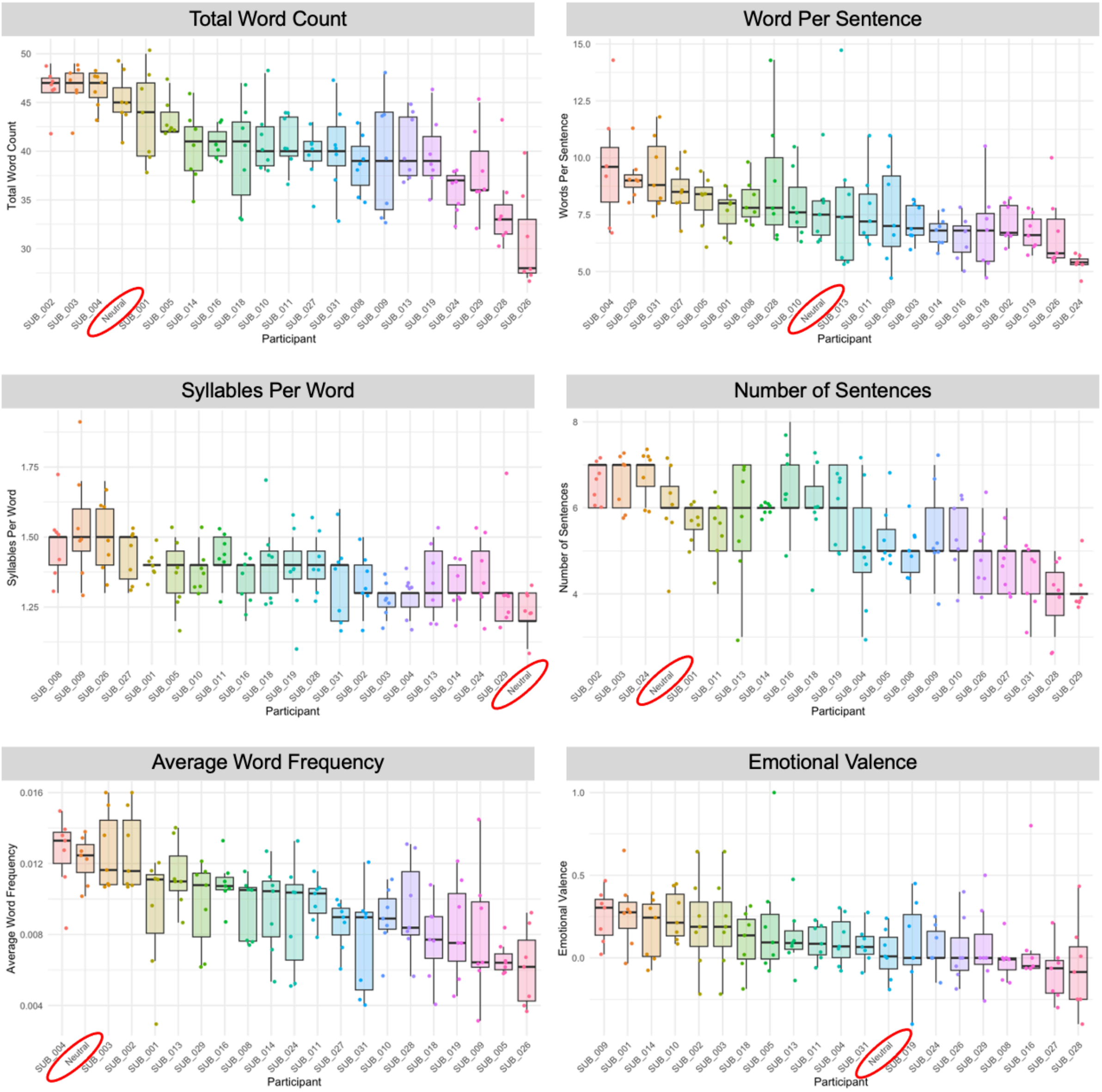

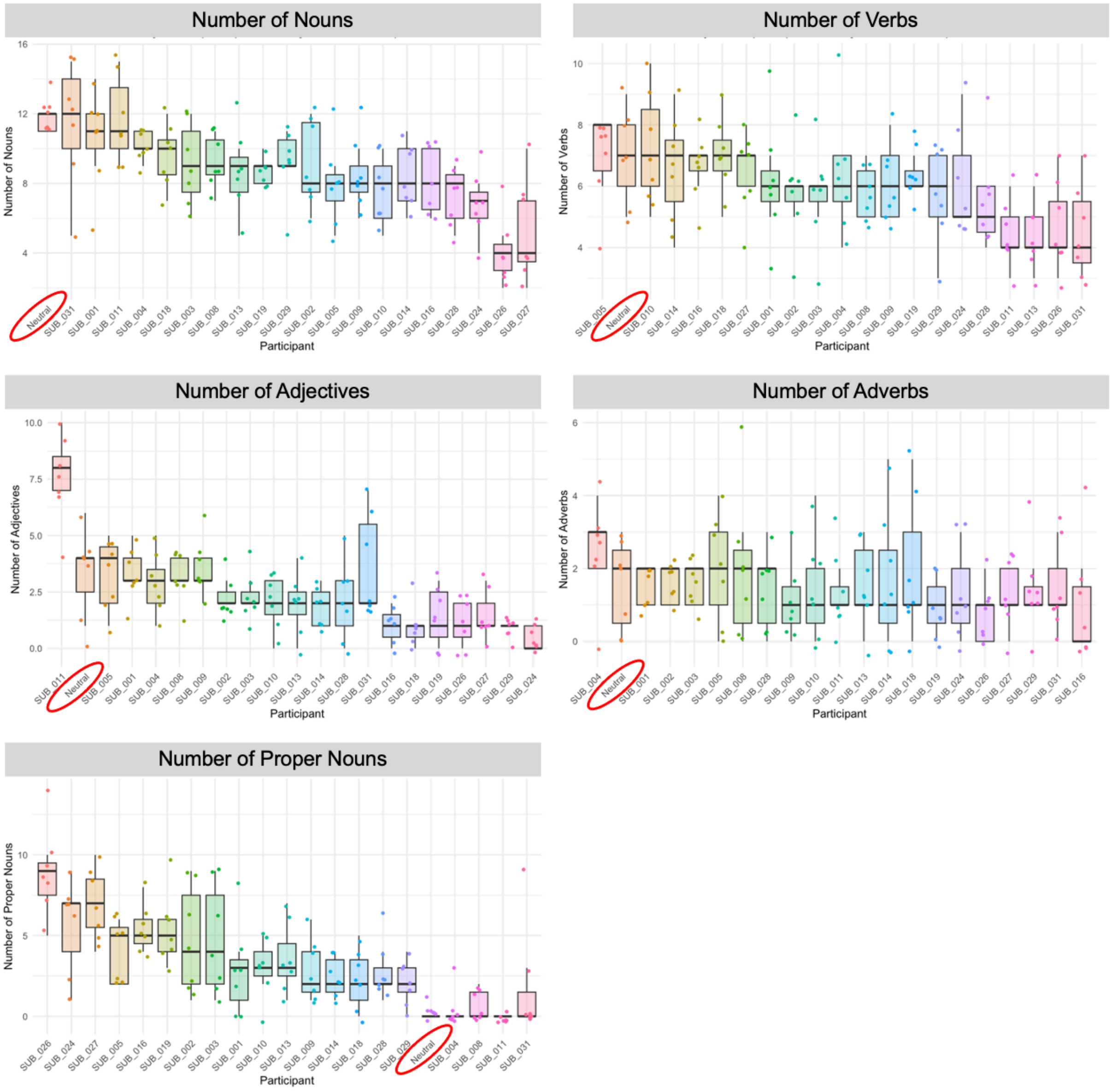


*Box plots show linguistic and paralinguistic features by participant, with each narrative represented by a dot. Plots are ordered by median values. The neutral condition is circled in red for reference.*

***Supplementary Table 1: Linguistic and paralinguistic features for individual participants' stimuli***

| Subject | Variable | Mean | Standard Deviation | Wilcoxon Statistic | P-value |
|---|---|---|---|---|---|
| Neutral | Total Word Count | 45.14 | 2.67 | NA | NA |
| SUB_001 | Total Word Count | 43.57 | 4.69 | 19.5 | 0.563 |
| SUB_002 | Total Word Count | 46.43 | 2.23 | 33 | 0.303 |
| SUB_003 | Total Word Count | 46.57 | 2.30 | 33.5 | 0.273 |
| SUB_004 | Total Word Count | 46.43 | 1.90 | 31.5 | 0.398 |
| SUB_005 | Total Word Count | 43.29 | 1.98 | 14 | 0.194 |
| SUB_008 | Total Word Count | 38.71 | 3.09 | 2 | 0.005 |
| SUB_009 | Total Word Count | 39.43 | 6.02 | 9.5 | 0.060 |
| SUB_010 | Total Word Count | 41.14 | 3.58 | 7.5 | 0.034 |
| SUB_011 | Total Word Count | 41.00 | 2.71 | 5 | 0.014 |
| SUB_013 | Total Word Count | 40.43 | 3.46 | 7 | 0.028 |
| SUB_014 | Total Word Count | 40.43 | 3.69 | 7.5 | 0.034 |
| SUB_016 | Total Word Count | 40.86 | 1.68 | 3 | 0.007 |
| SUB_018 | Total Word Count | 39.71 | 5.35 | 8.5 | 0.046 |
| SUB_019 | Total Word Count | 39.71 | 3.73 | 6 | 0.021 |
| SUB_024 | Total Word Count | 35.86 | 2.27 | 0 | **0.002** |
| SUB_026 | Total Word Count | 30.86 | 4.95 | 0 | **0.002** |
| SUB_027 | Total Word Count | 39.57 | 2.88 | 2 | 0.005 |
| SUB_028 | Total Word Count | 34.00 | 4.40 | 1 | 0.003 |
| SUB_029 | Total Word Count | 37.86 | 4.34 | 5 | 0.014 |
| SUB_031 | Total Word Count | 40.29 | 4.54 | 7.5 | 0.034 |
| Neutral | Words Per Sentence | 7.74 | 1.62 | NA | NA |
| SUB_001 | Words Per Sentence | 7.66 | 0.93 | 27.5 | 0.748 |
| SUB_002 | Words Per Sentence | 7.13 | 0.85 | 20 | 0.608 |
| SUB_003 | Words Per Sentence | 7.16 | 0.84 | 21 | 0.701 |
| SUB_004 | Words Per Sentence | 9.66 | 2.61 | 37 | 0.125 |
| SUB_005 | Words Per Sentence | 8.10 | 1.15 | 33 | 0.305 |
| SUB_008 | Words Per Sentence | 8.09 | 0.99 | 32 | 0.371 |
| SUB_009 | Words Per Sentence | 7.61 | 2.26 | 23.5 | 0.949 |
| SUB_010 | Words Per Sentence | 7.96 | 1.54 | 25.5 | 0.949 |
| SUB_011 | Words Per Sentence | 7.77 | 1.69 | 24 | 1.000 |
| SUB_013 | Words Per Sentence | 7.97 | 3.32 | 22 | 0.798 |
| SUB_014 | Words Per Sentence | 6.73 | 0.64 | 13.5 | 0.177 |
| SUB_016 | Words Per Sentence | 6.47 | 0.96 | 12 | 0.124 |
| SUB_018 | Words Per Sentence | 6.86 | 1.96 | 15.5 | 0.277 |
| SUB_019 | Words Per Sentence | 6.71 | 0.79 | 13 | 0.160 |
| SUB_024 | Words Per Sentence | 5.36 | 0.39 | 0 | **0.002** |
| SUB_026 | Words Per Sentence | 6.71 | 1.68 | 12.5 | 0.141 |
| SUB_027 | Words Per Sentence | 8.53 | 1.13 | 35.5 | 0.177 |
| SUB_028 | Words Per Sentence | 8.94 | 2.84 | 30.5 | 0.481 |
| SUB_029 | Words Per Sentence | 9.17 | 1.06 | 41.5 | 0.034 |
| SUB_031 | Words Per Sentence | 9.31 | 1.65 | 38.5 | 0.084 |
| Neutral | Syllables Per Word | 1.23 | 0.08 | NA | NA |
| SUB_001 | Syllables Per Word | 1.40 | 0.06 | 47.5 | 0.003 |

| | | | | | |
|---|---|---|---|---|---|
| SUB_002 | Syllables Per Word | 1.34 | 0.10 | 40 | 0.043 |
| SUB_003 | Syllables Per Word | 1.29 | 0.07 | 34 | 0.208 |
| SUB_004 | Syllables Per Word | 1.29 | 0.07 | 34 | 0.208 |
| SUB_005 | Syllables Per Word | 1.37 | 0.11 | 41.5 | 0.029 |
| SUB_008 | Syllables Per Word | 1.47 | 0.13 | 47.5 | 0.003 |
| SUB_009 | Syllables Per Word | 1.54 | 0.20 | 47.5 | 0.003 |
| SUB_010 | Syllables Per Word | 1.37 | 0.08 | 44.5 | 0.009 |
| SUB_011 | Syllables Per Word | 1.43 | 0.08 | 47.5 | 0.003 |
| SUB_013 | Syllables Per Word | 1.34 | 0.13 | 37 | 0.108 |
| SUB_014 | Syllables Per Word | 1.33 | 0.08 | 40 | 0.042 |
| SUB_016 | Syllables Per Word | 1.34 | 0.08 | 41.5 | 0.027 |
| SUB_018 | Syllables Per Word | 1.41 | 0.15 | 44.5 | 0.009 |
| SUB_019 | Syllables Per Word | 1.39 | 0.16 | 41 | 0.037 |
| SUB_024 | Syllables Per Word | 1.36 | 0.11 | 40 | 0.043 |
| SUB_026 | Syllables Per Word | 1.50 | 0.14 | 47.5 | 0.003 |
| SUB_027 | Syllables Per Word | 1.43 | 0.10 | 46 | 0.005 |
| SUB_028 | Syllables Per Word | 1.41 | 0.11 | 46 | 0.006 |
| SUB_029 | Syllables Per Word | 1.31 | 0.18 | 31 | 0.405 |
| SUB_031 | Syllables Per Word | 1.34 | 0.15 | 35.5 | 0.159 |
| Neutral | Number of Sentences | 6.00 | 1.00 | NA | NA |
| SUB_001 | Number of Sentences | 5.71 | 0.49 | 17 | 0.296 |
| SUB_002 | Number of Sentences | 6.57 | 0.53 | 33 | 0.253 |
| SUB_003 | Number of Sentences | 6.57 | 0.53 | 33 | 0.253 |
| SUB_004 | Number of Sentences | 5.14 | 1.46 | 15.5 | 0.264 |
| SUB_005 | Number of Sentences | 5.43 | 0.79 | 14 | 0.179 |
| SUB_008 | Number of Sentences | 4.86 | 0.69 | 8 | 0.033 |
| SUB_009 | Number of Sentences | 5.43 | 1.13 | 16.5 | 0.321 |
| SUB_010 | Number of Sentences | 5.29 | 0.76 | 12.5 | 0.114 |
| SUB_011 | Number of Sentences | 5.43 | 0.79 | 14.5 | 0.177 |
| SUB_013 | Number of Sentences | 5.71 | 1.50 | 23 | 0.893 |
| SUB_014 | Number of Sentences | 6.00 | 0.00 | 21 | 0.593 |
| SUB_016 | Number of Sentences | 6.43 | 0.98 | 29 | 0.580 |
| SUB_018 | Number of Sentences | 6.00 | 1.00 | 24.5 | 1.000 |
| SUB_019 | Number of Sentences | 6.00 | 1.00 | 24 | 1.000 |
| SUB_024 | Number of Sentences | 6.71 | 0.49 | 36 | 0.116 |
| SUB_026 | Number of Sentences | 4.71 | 0.76 | 7.5 | 0.028 |
| SUB_027 | Number of Sentences | 4.71 | 0.76 | 7.5 | 0.028 |
| SUB_028 | Number of Sentences | 4.00 | 0.82 | 3.5 | 0.007 |
| SUB_029 | Number of Sentences | 4.14 | 0.38 | 4 | 0.006 |
| SUB_031 | Number of Sentences | 4.43 | 0.79 | 5 | 0.013 |
| Neutral | Average Word Frequency | 0.0122 | 0.0013 | NA | NA |
| SUB_001 | Average Word Frequency | 0.0093 | 0.0034 | 8 | 0.041 |
| SUB_002 | Average Word Frequency | 0.0127 | 0.0023 | 27 | 0.798 |
| SUB_003 | Average Word Frequency | 0.0127 | 0.0023 | 27 | 0.798 |
| SUB_004 | Average Word Frequency | 0.0126 | 0.0022 | 32 | 0.371 |
| SUB_005 | Average Word Frequency | 0.0066 | 0.0009 | 0 | **0.002** |
| SUB_008 | Average Word Frequency | 0.0094 | 0.0018 | 6 | 0.021 |
| SUB_009 | Average Word Frequency | 0.0080 | 0.0037 | 8 | 0.041 |
| SUB_010 | Average Word Frequency | 0.0089 | 0.0018 | 3 | 0.007 |

| | | | | | |
|---|---|---|---|---|---|
| SUB_011 | Average Word Frequency | 0.0099 | 0.0013 | 5 | 0.015 |
| SUB_013 | Average Word Frequency | 0.0114 | 0.0019 | 19 | 0.523 |
| SUB_014 | Average Word Frequency | 0.0095 | 0.0026 | 9 | 0.055 |
| SUB_016 | Average Word Frequency | 0.0109 | 0.0014 | 12 | 0.125 |
| SUB_018 | Average Word Frequency | 0.0077 | 0.0023 | 2 | 0.005 |
| SUB_019 | Average Word Frequency | 0.0083 | 0.0028 | 4 | 0.011 |
| SUB_024 | Average Word Frequency | 0.0091 | 0.0031 | 9 | 0.055 |
| SUB_026 | Average Word Frequency | 0.0061 | 0.0022 | 0 | **0.002** |
| SUB_027 | Average Word Frequency | 0.0086 | 0.0014 | 0 | **0.002** |
| SUB_028 | Average Word Frequency | 0.0095 | 0.0028 | 11 | 0.097 |
| SUB_029 | Average Word Frequency | 0.0097 | 0.0025 | 8 | 0.041 |
| SUB_031 | Average Word Frequency | 0.0076 | 0.0030 | 2 | 0.005 |
| Neutral | Emotional Valence | 0.0253 | 0.1543 | NA | NA |
| SUB_001 | Emotional Valence | 0.2748 | 0.2168 | 42 | 0.030 |
| SUB_002 | Emotional Valence | 0.2040 | 0.2804 | 34 | 0.250 |
| SUB_003 | Emotional Valence | 0.2040 | 0.2804 | 34 | 0.250 |
| SUB_004 | Emotional Valence | 0.1051 | 0.1499 | 31 | 0.443 |
| SUB_005 | Emotional Valence | 0.2186 | 0.3725 | 33 | 0.307 |
| SUB_008 | Emotional Valence | -0.0133 | 0.1173 | 17 | 0.369 |
| SUB_009 | Emotional Valence | 0.2538 | 0.1596 | 43 | 0.021 |
| SUB_010 | Emotional Valence | 0.2549 | 0.1515 | 42 | 0.030 |
| SUB_011 | Emotional Valence | 0.0952 | 0.1087 | 32 | 0.371 |
| SUB_013 | Emotional Valence | 0.1229 | 0.1887 | 32 | 0.371 |
| SUB_014 | Emotional Valence | 0.1754 | 0.1898 | 36 | 0.160 |
| SUB_016 | Emotional Valence | 0.0865 | 0.3173 | 21.5 | 0.749 |
| SUB_018 | Emotional Valence | 0.1024 | 0.1788 | 32 | 0.371 |
| SUB_019 | Emotional Valence | 0.0703 | 0.2837 | 27 | 0.797 |
| SUB_024 | Emotional Valence | 0.0603 | 0.1381 | 26.5 | 0.846 |
| SUB_026 | Emotional Valence | 0.0438 | 0.2056 | 24 | 1.000 |
| SUB_027 | Emotional Valence | -0.0865 | 0.1724 | 12.5 | 0.141 |
| SUB_028 | Emotional Valence | -0.0595 | 0.2802 | 18 | 0.443 |
| SUB_029 | Emotional Valence | 0.0640 | 0.2505 | 23.5 | 0.948 |
| SUB_031 | Emotional Valence | 0.0795 | 0.1155 | 29.5 | 0.565 |
| Neutral | Number of Nouns | 11.86 | 1.07 | NA | NA |
| SUB_001 | Number of Nouns | 10.57 | 2.88 | 18.5 | 0.461 |
| SUB_002 | Number of Nouns | 9.14 | 2.48 | 10.5 | 0.074 |
| SUB_003 | Number of Nouns | 9.14 | 2.34 | 9 | 0.049 |
| SUB_004 | Number of Nouns | 10.29 | 0.76 | 4.5 | 0.009 |
| SUB_005 | Number of Nouns | 8.00 | 2.24 | 4.5 | 0.011 |
| SUB_008 | Number of Nouns | 9.29 | 1.50 | 3 | 0.006 |
| SUB_009 | Number of Nouns | 8.43 | 1.99 | 4.5 | 0.011 |
| SUB_010 | Number of Nouns | 7.57 | 1.99 | 0 | **0.002** |
| SUB_011 | Number of Nouns | 11.71 | 2.50 | 21.5 | 0.740 |
| SUB_013 | Number of Nouns | 8.71 | 2.50 | 6 | 0.020 |
| SUB_014 | Number of Nouns | 8.43 | 1.90 | 1.5 | 0.003 |
| SUB_016 | Number of Nouns | 8.14 | 1.86 | 0 | **0.002** |
| SUB_018 | Number of Nouns | 9.57 | 1.72 | 6 | 0.019 |
| SUB_019 | Number of Nouns | 8.71 | 0.76 | 0 | **0.002** |
| SUB_024 | Number of Nouns | 6.86 | 1.86 | 0 | **0.002** |

| | | | | | |
|---|---|---|---|---|---|
| SUB_026 | Number of Nouns | 4.14 | 1.95 | 0 | **0.002** |
| SUB_027 | Number of Nouns | 5.29 | 2.81 | 0 | **0.002** |
| SUB_028 | Number of Nouns | 7.29 | 1.60 | 0 | **0.002** |
| SUB_029 | Number of Nouns | 9.14 | 2.04 | 3 | 0.006 |
| SUB_031 | Number of Nouns | 11.43 | 3.55 | 26 | 0.896 |
| Neutral | Number of Verbs | 7.00 | 1.53 | NA | NA |
| SUB_001 | Number of Verbs | 6.14 | 2.12 | 17 | 0.364 |
| SUB_002 | Number of Verbs | 5.71 | 1.50 | 14 | 0.192 |
| SUB_003 | Number of Verbs | 5.71 | 1.50 | 14 | 0.192 |
| SUB_004 | Number of Verbs | 6.43 | 1.90 | 18 | 0.435 |
| SUB_005 | Number of Verbs | 7.00 | 1.53 | 25 | 1.000 |
| SUB_008 | Number of Verbs | 5.86 | 0.90 | 13 | 0.145 |
| SUB_009 | Number of Verbs | 6.14 | 1.21 | 16 | 0.288 |
| SUB_010 | Number of Verbs | 7.29 | 1.80 | 26.5 | 0.846 |
| SUB_011 | Number of Verbs | 4.43 | 0.98 | 4 | 0.009 |
| SUB_013 | Number of Verbs | 4.43 | 0.98 | 4 | 0.009 |
| SUB_014 | Number of Verbs | 6.57 | 1.72 | 20.5 | 0.648 |
| SUB_016 | Number of Verbs | 6.71 | 0.95 | 20 | 0.591 |
| SUB_018 | Number of Verbs | 7.00 | 1.29 | 23.5 | 0.947 |
| SUB_019 | Number of Verbs | 6.29 | 0.95 | 17 | 0.359 |
| SUB_024 | Number of Verbs | 6.14 | 1.68 | 17.5 | 0.384 |
| SUB_026 | Number of Verbs | 4.71 | 1.38 | 6 | 0.020 |
| SUB_027 | Number of Verbs | 6.43 | 1.27 | 18 | 0.430 |
| SUB_028 | Number of Verbs | 5.57 | 1.72 | 12.5 | 0.135 |
| SUB_029 | Number of Verbs | 5.71 | 1.50 | 13 | 0.145 |
| SUB_031 | Number of Verbs | 4.57 | 1.51 | 6 | 0.020 |
| Neutral | Number of Adjectives | 3.29 | 2.06 | NA | NA |
| SUB_001 | Number of Adjectives | 3.29 | 1.25 | 21.5 | 0.738 |
| SUB_002 | Number of Adjectives | 2.29 | 0.95 | 15.5 | 0.261 |
| SUB_003 | Number of Adjectives | 2.29 | 0.95 | 15.5 | 0.261 |
| SUB_004 | Number of Adjectives | 2.86 | 1.35 | 19.5 | 0.555 |
| SUB_005 | Number of Adjectives | 3.29 | 1.60 | 25.5 | 0.947 |
| SUB_008 | Number of Adjectives | 3.14 | 1.07 | 19.5 | 0.537 |
| SUB_009 | Number of Adjectives | 3.57 | 1.27 | 22.5 | 0.841 |
| SUB_010 | Number of Adjectives | 2.00 | 1.15 | 12 | 0.118 |
| SUB_011 | Number of Adjectives | 7.57 | 1.90 | 46 | 0.006 |
| SUB_013 | Number of Adjectives | 2.00 | 1.29 | 14 | 0.188 |
| SUB_014 | Number of Adjectives | 1.86 | 0.90 | 12.5 | 0.132 |
| SUB_016 | Number of Adjectives | 1.00 | 0.82 | 9.5 | 0.057 |
| SUB_018 | Number of Adjectives | 1.00 | 1.00 | 9 | 0.046 |
| SUB_019 | Number of Adjectives | 1.43 | 1.27 | 10 | 0.068 |
| SUB_024 | Number of Adjectives | 0.43 | 0.53 | 6.5 | 0.019 |
| SUB_026 | Number of Adjectives | 1.14 | 0.90 | 10 | 0.067 |
| SUB_027 | Number of Adjectives | 1.43 | 1.13 | 10.5 | 0.074 |
| SUB_028 | Number of Adjectives | 2.14 | 1.77 | 15 | 0.242 |
| SUB_029 | Number of Adjectives | 0.86 | 0.38 | 9.5 | 0.045 |
| SUB_031 | Number of Adjectives | 3.71 | 2.21 | 27.5 | 0.744 |
| Neutral | Number of Adverbs | 1.57 | 1.27 | NA | NA |
| SUB_001 | Number of Adverbs | 1.57 | 0.53 | 23.5 | 0.946 |

| | | | | | |
|---|---|---|---|---|---|
| SUB_002 | Number of Adverbs | 1.57 | 0.53 | 23.5 | 0.946 |
| SUB_003 | Number of Adverbs | 1.57 | 0.53 | 23.5 | 0.946 |
| SUB_004 | Number of Adverbs | 2.43 | 1.27 | 34 | 0.232 |
| SUB_005 | Number of Adverbs | 2.00 | 1.53 | 29 | 0.597 |
| SUB_008 | Number of Adverbs | 2.00 | 2.08 | 25.5 | 0.948 |
| SUB_009 | Number of Adverbs | 1.14 | 1.07 | 19.5 | 0.553 |
| SUB_010 | Number of Adverbs | 1.43 | 1.40 | 22 | 0.793 |
| SUB_011 | Number of Adverbs | 1.29 | 0.95 | 21 | 0.691 |
| SUB_013 | Number of Adverbs | 1.57 | 1.13 | 24.5 | 1.000 |
| SUB_014 | Number of Adverbs | 1.71 | 1.80 | 24 | 1.000 |
| SUB_016 | Number of Adverbs | 1.00 | 1.53 | 17.5 | 0.384 |
| SUB_018 | Number of Adverbs | 2.00 | 1.83 | 26.5 | 0.845 |
| SUB_019 | Number of Adverbs | 1.00 | 0.82 | 17.5 | 0.390 |
| SUB_024 | Number of Adverbs | 1.29 | 1.25 | 21.5 | 0.741 |
| SUB_026 | Number of Adverbs | 0.71 | 0.76 | 14.5 | 0.206 |
| SUB_027 | Number of Adverbs | 1.29 | 0.76 | 20.5 | 0.642 |
| SUB_028 | Number of Adverbs | 1.43 | 1.13 | 22.5 | 0.842 |
| SUB_029 | Number of Adverbs | 1.43 | 1.27 | 22 | 0.792 |
| SUB_031 | Number of Adverbs | 1.43 | 1.13 | 23 | 0.894 |
| Neutral | Number of Proper Nouns | 0.14 | 0.38 | NA | NA |
| SUB_001 | Number of Proper Nouns | 2.86 | 2.73 | 41 | 0.023 |
| SUB_002 | Number of Proper Nouns | 4.71 | 3.35 | 48.5 | **0.002** |
| SUB_003 | Number of Proper Nouns | 4.71 | 3.35 | 48.5 | **0.002** |
| SUB_004 | Number of Proper Nouns | 0.43 | 1.13 | 25 | 1.000 |
| SUB_005 | Number of Proper Nouns | 4.00 | 1.91 | 49 | **0.001** |
| SUB_008 | Number of Proper Nouns | 0.71 | 0.95 | 32.5 | 0.228 |
| SUB_009 | Number of Proper Nouns | 2.86 | 1.86 | 48 | **0.002** |
| SUB_010 | Number of Proper Nouns | 3.00 | 1.73 | 45 | 0.006 |
| SUB_011 | Number of Proper Nouns | 0.00 | 0.00 | 21 | 0.391 |
| SUB_013 | Number of Proper Nouns | 3.57 | 2.15 | 48.5 | **0.002** |
| SUB_014 | Number of Proper Nouns | 2.43 | 1.27 | 48 | **0.002** |
| SUB_016 | Number of Proper Nouns | 5.43 | 1.40 | 49 | **0.001** |
| SUB_018 | Number of Proper Nouns | 2.29 | 1.89 | 41 | 0.023 |
| SUB_019 | Number of Proper Nouns | 5.43 | 2.30 | 49 | **0.001** |
| SUB_024 | Number of Proper Nouns | 5.57 | 2.94 | 48.5 | **0.002** |
| SUB_026 | Number of Proper Nouns | 8.86 | 2.79 | 49 | **0.001** |
| SUB_027 | Number of Proper Nouns | 7.00 | 2.16 | 49 | **0.001** |
| SUB_028 | Number of Proper Nouns | 2.71 | 1.70 | 48.5 | **0.002** |
| SUB_029 | Number of Proper Nouns | 2.14 | 1.35 | 44.5 | 0.008 |
| SUB_031 | Number of Proper Nouns | 1.71 | 3.40 | 29 | 0.477 |
| Neutral | Number of Nouns (combined) | 12.00 | 1.41 | NA | NA |
| SUB_001 | Number of Nouns (combined) | 13.43 | 1.40 | 40 | 0.048 |
| SUB_002 | Number of Nouns (combined) | 13.86 | 1.35 | 42 | 0.027 |
| SUB_003 | Number of Nouns (combined) | 13.86 | 1.57 | 41 | 0.035 |
| SUB_004 | Number of Nouns (combined) | 10.71 | 1.25 | 10.5 | 0.071 |
| SUB_005 | Number of Nouns (combined) | 12.00 | 2.08 | 23 | 0.896 |
| SUB_008 | Number of Nouns (combined) | 10.00 | 2.00 | 9 | 0.049 |
| SUB_009 | Number of Nouns (combined) | 11.29 | 1.38 | 21 | 0.686 |
| SUB_010 | Number of Nouns (combined) | 10.57 | 2.44 | 11 | 0.082 |

| | | | | | |
|---|---|---|---|---|---|
| SUB_011 | Number of Nouns (combined) | 11.71 | 2.50 | 20.5 | 0.642 |
| SUB_013 | Number of Nouns (combined) | 12.29 | 1.89 | 27.5 | 0.742 |
| SUB_014 | Number of Nouns (combined) | 10.86 | 2.54 | 19.5 | 0.552 |
| SUB_016 | Number of Nouns (combined) | 13.57 | 2.07 | 36.5 | 0.134 |
| SUB_018 | Number of Nouns (combined) | 11.86 | 2.67 | 25.5 | 0.948 |
| SUB_019 | Number of Nouns (combined) | 14.14 | 2.12 | 39 | 0.068 |
| SUB_024 | Number of Nouns (combined) | 12.43 | 1.99 | 30.5 | 0.471 |
| SUB_026 | Number of Nouns (combined) | 13.00 | 2.58 | 32 | 0.360 |
| SUB_027 | Number of Nouns (combined) | 12.29 | 2.29 | 25 | 1.000 |
| SUB_028 | Number of Nouns (combined) | 10.00 | 1.00 | 3 | 0.005 |
| SUB_029 | Number of Nouns (combined) | 11.29 | 1.38 | 21 | 0.686 |
| SUB_031 | Number of Nouns (combined) | 13.14 | 1.57 | 35.5 | 0.163 |

*Non-parametric pairwise comparisons were conducted using Wilcoxon rank-sum tests for each individual subjects' narratives vs. neutral narratives. Significant differences are bolded, using a Bonferroni correction for multiple comparisons (adjusted p<0.0025 for 20 comparisons).*

### PERSONALIZED VIDEOS TASK, ANALYSES, AND DISCUSSION

**Task design**. Participants were asked to passively watch video clips (accompanied by audio) in a block-design paradigm (mirroring the personalized narratives task). The task consisted of two conditions: INTEREST and NEUTRAL. In the INTEREST condition, participants watched the 16-second video clips derived directly from the videos they provided (see **Personalized Stimuli Creation** in Main Text). In the NEUTRAL condition, participants watched 16-second non-personalized videos of nature scenes. Each video was followed by an inter-stimulus rest block of 5 seconds cued by a grey fixation cross (total of 14 videos across two conditions and 15 rest blocks). To confirm that children were attending to the task without imposing significant physical or cognitive demands, we included a low-demand attentional check following each video. An image of a panda appeared on the screen directly after each video for 1.5 seconds, followed by a grey fixation and screen for 0.5 seconds. Children were instructed at the beginning of the study to press a button using their pointer finger via an MRI-compatible button box every time they saw a picture of a panda. Task order was fixed across participants in an [ABAB…] pattern: INTEREST then NEUTRAL.

**Analyses.** Preprocessing for fMRI data arising from the personalized videos task was identical to that described in methods. Whole-brain analyses were conducted to examine the effect of personal interest videos on the brain. Group-level modeling was performed using SPM12. First-level maps from several contrasts of interest (NEUTRAL, INTEREST, INTEREST > NEUTRAL) were brought up to a second-level analyses wherein one-sample t-tests were used to determine regions for which activation in the condition of interest was greater than baseline. Group maps were thresholded at an uncorrected voxel $p < 0.001$, with a cluster correction for multiple comparisons (FWE < 0.05).

**Results.** As for the personal interest narrative task, personal interest videos resulted in greater activation in several brain regions when compared to neutral videos. However, unlike the narrative task, many regions showing increased activation were associated with visual rather than language processing including bilateral primary visual cortex, bilateral lateral occipital cortices (LOC), bilateral inferior temporal cortices and fusiform gyrus, bilateral posterior parietal cortex, and right frontal regions.

**Discussion**. The personalized videos task was designed to be highly naturalistic to reflect children's patterns of brain activation as they authentically engaged with their interest. We solicited timestamps for the videos from families to capture each child's favorite or most salient part of the videos. Unsurprisingly, the INTEREST video clips varied immensely across many dimensions between participants. Some clips had language, some had music; some had humans, some had animals; some had large amounts of movement and action, some had very little movement. The NEUTRAL condition featured nature scenes with no humans, no language, and limited movement (e.g., grass blowing in the wind; snow falling in the forest). Given the variability in the videos, and the many differences between the INTEREST and NEUTRAL conditions on the video task, it is difficult to interpret any specific differences in activation elicited between the conditions. However, it is notable that brain regions engaged by INTEREST>NEUTRAL on the video task were not the same as those engaged by the personalized language task. While the language task elicited responses in language regions, the videos task elicited responses in primarily visual processing regions (activation in these regions was not observed in for INTEREST>NEUTRAL on the narrative task). Thus, while not the focus of the current paper, the group whole-brain responses on the videos task show that the INTEREST>NEUTRAL contrast in the language task is not identifying a modality-independent set of regions always engaged by any type of interesting materials, otherwise the same network would have been observed for this contrast in the videos task. Similar to what we observed in the language task, an overlapping set of regions are engaged in both the interest and neutral conditions, but interest elicits higher responses in these regions. This further supports the interpretation of a *modulating* effect of interest on modality-specific brain networks.

***Supplementary Figure 3: Higher responses to personally interesting videos than generic (nature) videos***

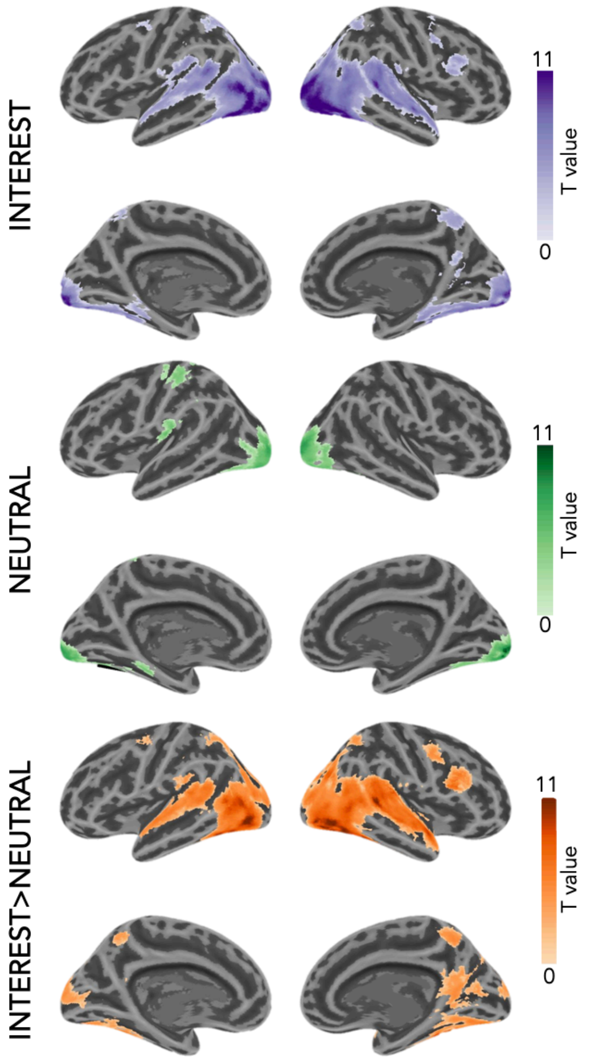


*Whole-brain activation maps for the INTEREST, NEUTRAL, and INTEREST>NEUTRAL contrasts from the personalized videos task (threshold: p<0.001, FWE corrected at p<0.05). Color bars reflect t-values.*
